# The molecular basis for functional divergence of duplicated SOX factors controlling endoderm formation and left-right patterning in zebrafish

**DOI:** 10.1101/2024.02.06.579092

**Authors:** Simaran Johal, Randa Elsayed, Kristen A. Panfilio, Andrew C. Nelson

**Author notes:** Corresponding author (ACN).

## Abstract

Endoderm, one of three primary germ layers of vertebrate embryos, makes major contributions to the respiratory and gastrointestinal tracts and associated organs, including liver and pancreas. In mammals, the transcription factor *SOX17* is vital for endoderm organ formation and can induce endoderm progenitor identity. Duplication of ancestral *sox17* in the teleost lineage produced the paralogues *sox32* and *sox17* in zebrafish. Sox32 is required for specification of endoderm and progenitors of the left-right organiser (Kupffer’s Vesicle, KV), with Sox17 a downstream target of Sox32 that is implicated in further KV development. Phenotypic evidence therefore suggests functional similarities between zebrafish Sox32 and Sox17 and mammalian SOX17. Here, we directly compare these orthologues and paralogues, using the early zebrafish embryo as a biological platform for functional testing. Our results indicate that, unlike Sox32, human SOX17 cannot induce endoderm specification in zebrafish. Furthermore, using hybrid protein functional analyses, we show that Sox32 specificity for the endoderm gene regulatory network is linked to evolutionary divergence in its DNA-binding HMG domain from its paralogue Sox17. Additionally, changes in the C-terminal regions of Sox32 and Sox17 underpin their differing target specificities. Finally, we establish that specific conserved peptides in the C-terminal domain are essential for the role of Sox17 in establishing correct organ asymmetry. Overall, our results illuminate the molecular basis for functional divergence of Sox32 and Sox17 in vertebrate endoderm development and left-right patterning, and the evolution of SoxF transcription factor function.

## Introduction

Sox family transcription factors (TFs) are evolutionarily conserved proteins with roles in cell fate decisions in a range of developmental processes (Sarkar and Hochedlinger 2013). Mutations in Sox factors have been linked to an array of developmental defects and congenital diseases in humans (Angelozzi and Lefebvre 2019). Sox TFs belong to the superfamily of High Mobility Group (HMG) domain containing TFs. Historically, membership of the Sox family has been dictated by having >50% conservation of the DNA-binding HMG domain with mouse SRY – the founding Sox TF (Gubbay, et al. 1990; Sinclair, et al. 1990; Pevny and Lovell-Badge 1997). Sox factors have been further characterised into subfamilies, whereby members of a subfamily exhibit 75-80% homology within their HMG domain (Bowles, et al. 2000; Wegner 2010). One subfamily is SoxF, which in mammals consists of SOX7, SOX17 and SOX18. In teleosts, SoxF is expanded to include Sox32. How has this additional TF been functionally integrated into SoxF developmental gene regulatory networks (GRNs)?

Most duplicated *sox* genes in the teleost lineage arose through a whole genome duplication event that occurred 226-316 million years ago (Voldoire, et al. 2017). An exception to this is *sox32,* which is hypothesised to have emerged through tandem duplication of *Sox17* (Voldoire, et al. 2017). While we and others have speculated that zebrafish Sox32 may function in similar processes to mammalian SOX17 (Lilly, et al. 2017; Figiel, et al. 2021), this has not been explored at the molecular level. Furthermore, it is unknown whether functions of the ancestral gene have been split among the two paralogues (sub-functionalisation) or whether they have functionally diverged post-duplication and acquired new roles (neo-functionalisation).

Sox32 is considered the master-regulator of endoderm identity in zebrafish. Loss-of-function mutations in *sox32* lead to a profound loss of endoderm progenitors and consequent absence of the respiratory tract and gut, and its associated organs including the liver and pancreas. Instead, presumptive endoderm cells differentiate into mesoderm (Alexander, et al. 1999; Dickmeis, et al. 2001). Conversely, *sox32* overexpression (OE) through mRNA injection during early development is sufficient to respecify presumptive mesodermal cells to endodermal fates, resulting in ectopic endoderm formation (Kikuchi, et al. 2001). Furthermore, transplantation studies show that cells from *sox32* mRNA-injected donor zebrafish embryos preferentially incorporate into developing endoderm tissues in wild type hosts, and they are capable of endoderm restoration in Sox32-deficient hosts (Kikuchi, et al. 2001; Aoki, et al. 2002; Stafford, et al. 2006).

Similarly, *Sox17* null mutant mice are embryonic lethal and show narrower expression domains of definitive endoderm markers in the embryonic gut, indicating that loss of Sox17 results in depletion of the definitive endoderm (Kanai-Azuma, et al. 2002). Furthermore, ectopic expression of SOX17 in mouse and human embryonic stem cells (ESCs) causes differentiation towards the extraembryonic and definitive endoderm fates, respectively (Séguin, et al. 2008; Niakan, et al. 2010; McDonald, et al. 2014). Thus, phenotypic evidence suggests functional similarity between SoxF subfamily members, with both mammalian SOX17 and zebrafish Sox32 inducing endoderm cell identity within their respective species. However, a molecular functional comparison is lacking.

In zebrafish, Sox32 is also required for correct specification of the Dorsal Forerunner Cells (DFCs) – progenitors of the left-right (LR) organiser, which is essential for correct positioning of endoderm-derived organs within the body cavity (Alexander, et al. 1999; Essner, et al. 2005). Specifically, the DFCs give rise to the zebrafish laterality organ, known as Kupffer’s Vesicle (KV) (Cooper and D’Amico 1996; Melby, et al. 1996; Essner, et al. 2005; Warga and Kane 2018). KV function involves cilia-mediated fluid flow in an anti-clockwise direction, which leads to unilateral induction in the left lateral plate mesoderm (LPM) of Nodal (Spaw in teleosts, (Long, et al. 2003; Montague, et al. 2018)). Defects in KV formation and cilia function lead to abnormalities in Nodal expression and consequent laterality defects affecting organs including the brain, heart, and pancreas (Essner, et al. 2005; Kramer-Zucker, et al. 2005). As a downstream target of its paralogue Sox32, Sox17 is implicated in KV function (Aamar and Dawid 2010). Similarly, in mice Sox17 is implicated in the establishment of LR patterning, both indirectly through an inability to maintain gut endoderm in *Sox17*^-/-^ mutants, and directly through defects in the morphogenesis of the mouse laterality organ (the node) (Kanai-Azuma, et al. 2002; Saund, et al. 2012). An outstanding question is which regions of the Sox17 protein mediate key functions in establishment of LR asymmetry and consequent organ patterning and placement.

Building on previous phenotypic observations, and to gain novel insights at the molecular level, here we examine whether SoxF factors exhibit functional conservation across species and identify functional domains within the zebrafish Sox32 and Sox17 proteins that are required for their respective developmental roles in endoderm specification and LR patterning. We performed molecular dissection and hybrid protein functional analyses of zebrafish Sox32 and Sox17 to draw functional comparisons with human SOX17 (hSOX17). We find that Sox32 shares limited functional similarities with hSOX17 or Sox17, with neither of the latter two able to induce endoderm specification in the zebrafish embryo. Sox17 and hSOX17 also differ from each other in terms of molecular capabilities. In detail, Sox32’s fundamental role in initiating the zebrafish endoderm GRN appears to be linked to evolutionary divergence in its HMG domain. We also show how changes in putative helical peptides within their C-terminal domains (CTDs) account for the zebrafish paralogues’ differing functional roles, including in cell fate specification and organ placement. Overall, our analyses reveal how divergence in specific protein domains underpins distinct TF target gene repertoires. We thus elucidate the SoxF molecular evolutionary changes that determine endoderm GRN specificity in the teleost and mammalian lineages.

## Results

### hSOX17 cannot induce endoderm identity in zebrafish embryos

Although hSOX17 and Sox32 can induce endoderm in humans and zebrafish, respectively, these orthologues’ molecular capabilities have not been tested in a directly comparable biological context. Also, it is unknown whether Sox17, downstream of Sox32, can induce endoderm gene expression directly. To address this, we injected equimolar quantities of mRNA encoding hSOX17, Sox32 or Sox17 into the zebrafish embryo at the one-cell stage, followed by RNA-seq analysis at 6 hours post-fertilisation (hpf), the stage by which endoderm has been specified in wild type embryos (Figure 1A).

**Figure 1:**
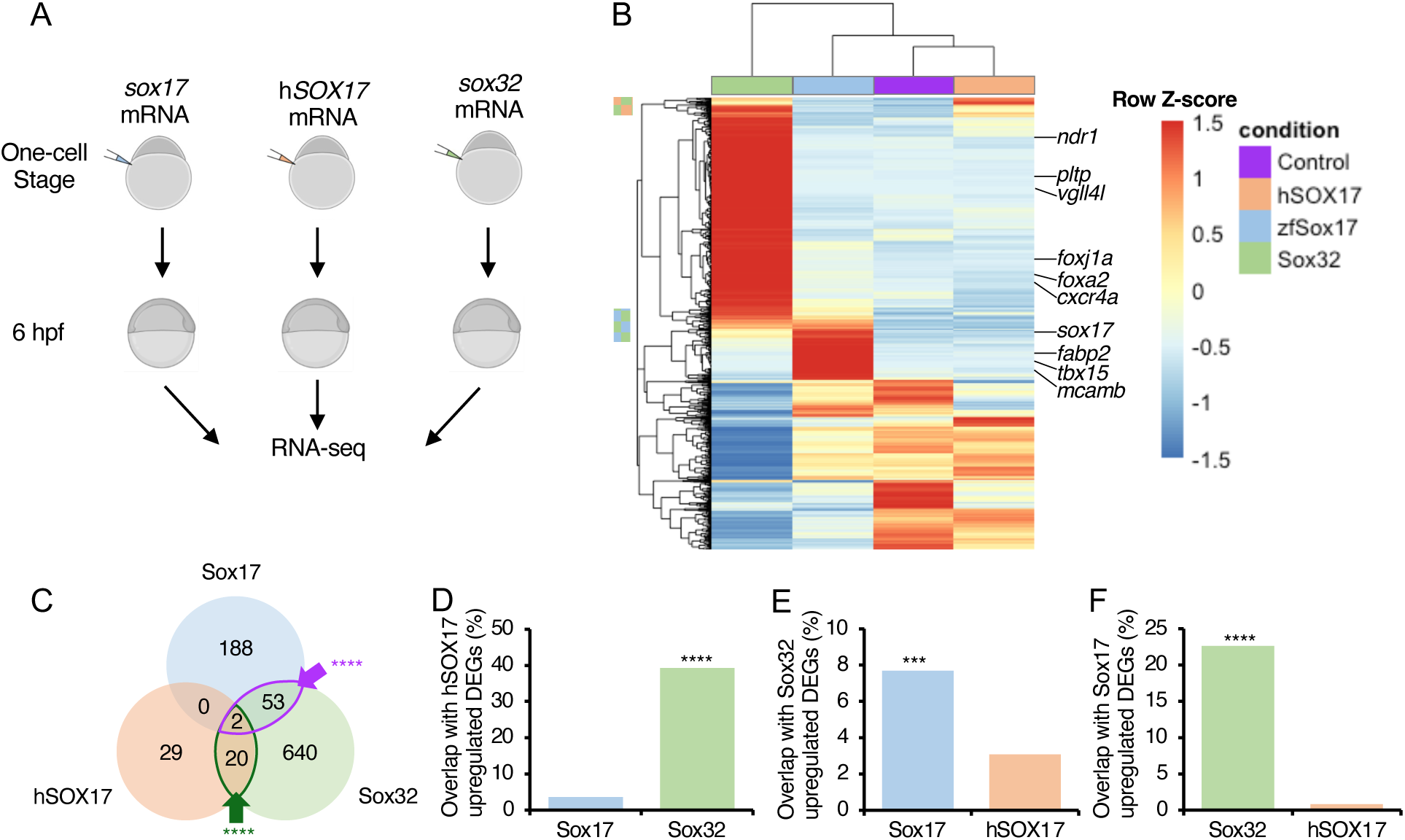
Sox32, Sox17 and hSOX17 are capable of inducing largely distinct gene sets in the early zebrafish embryo. **A)** Experimental schematic. **B)** Heatmap of all 1628 differentially expressed genes (DEGs, Supplementary Table S1-3), displayed as median counts from N=3 per condition. Colour-coded blocks to the left indicate genes induced by both hSOX17 and Sox32, and by Sox17 and Sox32. **C)** Venn diagram showing overlap of genes (Supplementary Table S4) significantly upregulated by each TF (P < 0.05). Overlaps significantly greater than expected according to Fisher’s Exact test are indicated. **** P < 0.0001. **D-F)** Bar charts indicating percentage overlap of upregulated DEGs to (D) hSOX17, (E) Sox32, and (F) Sox17 upregulated DEGs. Bars represent the percentage of genes upregulated by the factor on the y-axis also upregulated by the factor on the x-axis; note the TF-specific y-axis scale for each chart. Statistical tests to determine whether the overlap was significantly greater with one factor on the x-axis compared to the other were carried out using Fisher’s Exact test. *** P<0.001, **** P <0.0001.

Sox32 overexpression (OE) elicited widespread changes in gene expression compared to Sox17, with hSOX17 affecting relatively few genes (Figure 1B-C, Supplementary Figure S1, Supplementary Tables S1-5). We observed a small but significant overlap in upregulated transcripts between Sox32 and hSOX17, and Sox32 and Sox17, but not between hSOX17 and Sox17 (Figure 1C-F). Thus, Sox32 appears functionally more similar to both hSOX17 and Sox17 than hSOX17 and Sox17 are to each other in the context of the early zebrafish embryo. Overall, this suggests key differences in the functional capabilities of these three Sox factors.

To examine the pathways and processes influenced by OE of these TFs, we performed functional and anatomical annotation analysis of the differentially expressed genes (DEGs), including comparison to established cell identity markers from single-cell (sc)RNA-seq time course analysis of the early embryo (Wagner, et al. 2018). This revealed largely distinct results for each TF, further supporting the notion that they are functionally distinct in this context (Supplementary Figure S2-S3). As expected, transcripts upregulated by Sox32 OE were significantly enriched for terms associated with endoderm development (Supplementary Figure S3). This includes 6 hpf “nondorsal involuted anterior”, which contains endoderm progenitors, and 8 hpf endoderm. Transcripts induced by Sox32 also showed strong enrichment for 8 hpf DFC markers, suggesting that Sox32 is not only necessary (Alexander, et al. 1999) but also sufficient to drive DFC gene expression. In comparison, Sox17 OE results in a less profound effect on endoderm marker expression, with no significant effect on 6 hpf nondorsal involuted anterior marker induction but modest significant enrichment for endoderm makers at 8 hpf and for DFC markers, consistent with a role in KV formation and function (Supplementary Figure S3). Meanwhile, genes upregulated after hSOX17 ectopic expression did not yield any significantly enriched gene ontology terms, and downregulated genes do not pertain to endoderm but may be indicative of mis-regulation of Wnt signalling (Supplementary Figure S2).

Beyond endoderm development, analysis of transcripts upregulated by Sox17 revealed significant enrichment for terms associated with cardiovascular development, such as “Regulation of Sprouting Angiogenesis”, “Endothelial Tube Morphogenesis” and “Endocardium” (Supplementary Figure S2). This is consistent with *sox17* expression patterns during cardiovascular development in both zebrafish and mammals (Chung, et al. 2011; Saba, et al. 2019) (Supplementary Figure S4, Supplementary Video S1), and known *Sox17* loss-of-function phenotypes in mice vasculature and heart (Kanai-Azuma, et al. 2002; Saba, et al. 2019).

Thus, functional and anatomical enrichment analyses suggest Sox17 has different roles in early developmental processes compared to its paralogue Sox32, and hSOX17 cannot induce endoderm specification in zebrafish.

### Sox32 has a divergent HMG domain and longer C-terminal helix

To determine which structural motifs may explain the functional differences between Sox32, Sox17 and hSOX17, we analysed AlphaFold structural predictions and performed protein sequence alignments (Figure 2A-E). 3D structural models indicate a very high confidence prediction in the centre of all three proteins corresponding to the HMG domain. The HMG domains of Sox17 and hSOX17 are most similar, with 92% amino acid identity, while the Sox32 HMG domain differs from both Sox17 and hSOX17 with 67% and 69% identity, respectively (Figure 2D). Interestingly, 44% of amino acid differences between Sox32 and Sox17 correspond to changes in chemical properties that may influence the 3D folding of the protein, resulting in exposure of different residues to binding partners.

**Figure 2:**
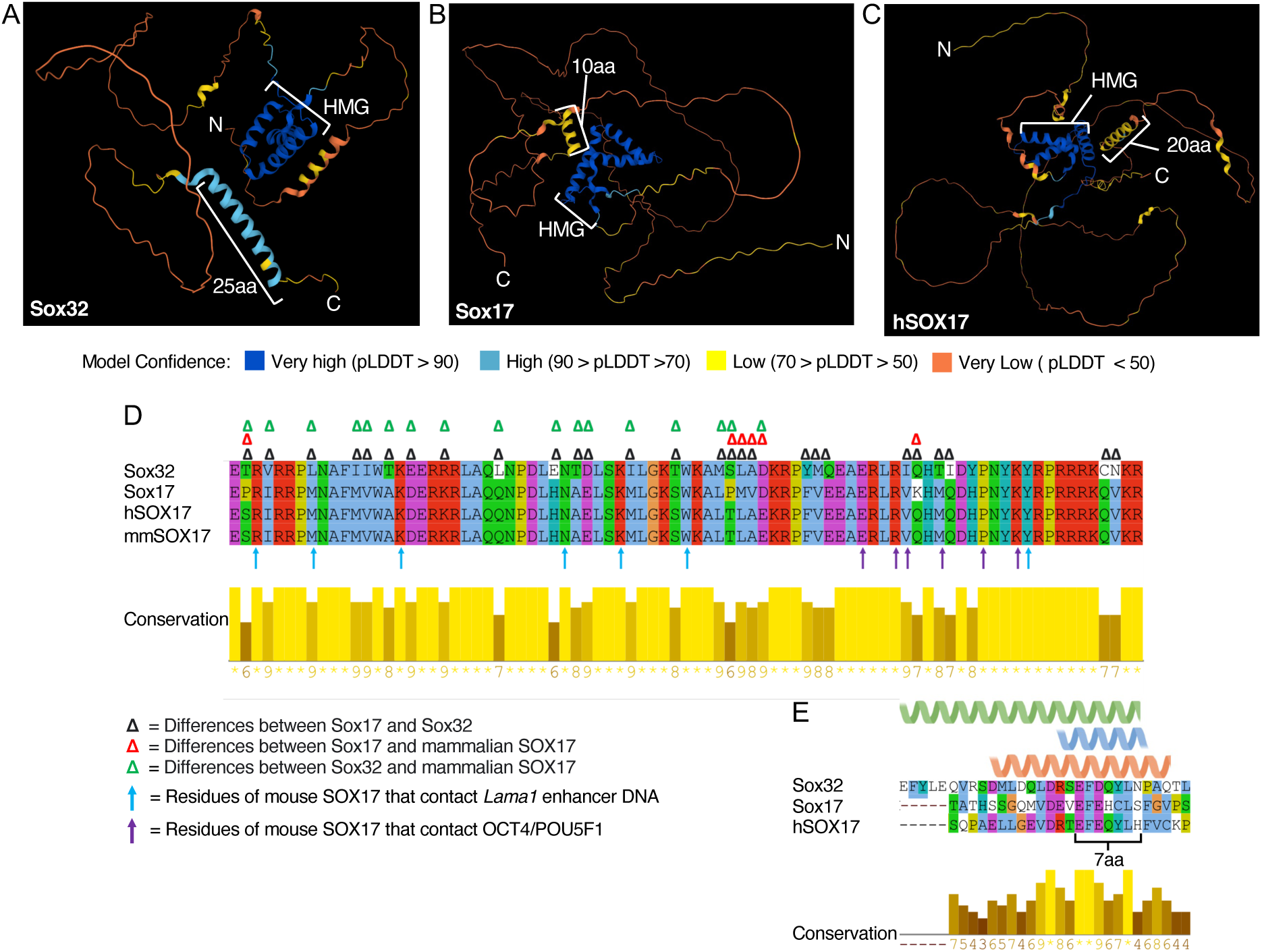
AlphaFold predicts C-terminal helical domains within Sox32, Sox17 and hSOX17. **A-C)** Alphafold models coloured by model confidence as shown in the key. DNA-binding HMG domains and C-terminal helices of different lengths are annotated. **(D)** Alignment of HMG domains of Sox32, Sox17, hSOX17 and *Mus musculus* (mm)Sox17. Alignment was carried out using Clustal Omega in Jalview, and colour coded based on conserved amino acid properties. Conservation levels calculated according to Analysis of Multiply Aligned Sequences (AMAS) displayed. Delta symbols denote differences in amino acids as per the key. Arrows indicating mapped amino acid contact sites between mouse SOX17 and the *Lama1* enhancer (Blue), and equivalent SOX2 sites and OCT4/POU5F1 protein (Purple). (Palasingam, et al. 2009; Ng, et al. 2012). **(E)** Alignment of CTD helical domain region for Sox32, Sox17 and hSOX17. Amino acids within Alphafold predicted helical domains annotated: Green = Sox32; blue = Sox17; peach = hSOX17. The 7-aa peptide previously shown to be required for Sox32 induction of endoderm fate (Zhao, et al. 2013) is also annotated.

Another notable structured region within Sox32 is a predicted 25-aa C-terminal helix (Figure 2A). C-terminal helices were also predicted in Sox17 (Figure 2B) and hSOX17 (Figure 2C), albeit the confidence of the prediction is lower and they are shorter (Figure 2E). Amino acid sequence alignments of the putative helices suggest the overall level of conservation is low, although they do encompass a seven amino acid peptide region previously recognised to be conserved throughout the SoxF subfamily, including in Sox7 and Sox18 (Kikuchi, et al. 2001; Sinner, et al. 2004) (Supplementary Figure S5). This 7-aa Sox32 peptide (EFDQYLN, Figure 2E) has previously been shown to be required for Sox32 induction of endoderm fate (Zhao, et al. 2013).

Overall, we conclude that the Sox32 HMG domain shows a higher degree of divergence from the common ancestor than Sox17 or hSOX17. Additionally, we identify putative CTD helices of differing lengths between these Sox factors that encompass a 7-aa peptide region of known importance in Sox32.

### Sox32 specificity for the endoderm GRN is driven by HMG domain divergence

Compared to its homologues, Sox32 has a unique property allowing it to induce the zebrafish endoderm GRN (Figure 1), and divergence in the DNA-binding HMG domain (Figure 2D) may confer this function. To examine this, we tested the ability of a hybrid protein (HMG Switch), consisting of the Sox32 N- and CTDs flanking the Sox17 HMG domain, to induce endoderm gene expression, compared to wild type Sox32 and Sox17 proteins and a Sox32 protein lacking the HMG domain (Sox32ΔHMG, Figure 3A). These proteins’ transcriptional effects were assayed by RNA-seq after early ectopic mRNA injection (Figure 3B).

**Figure 3:**
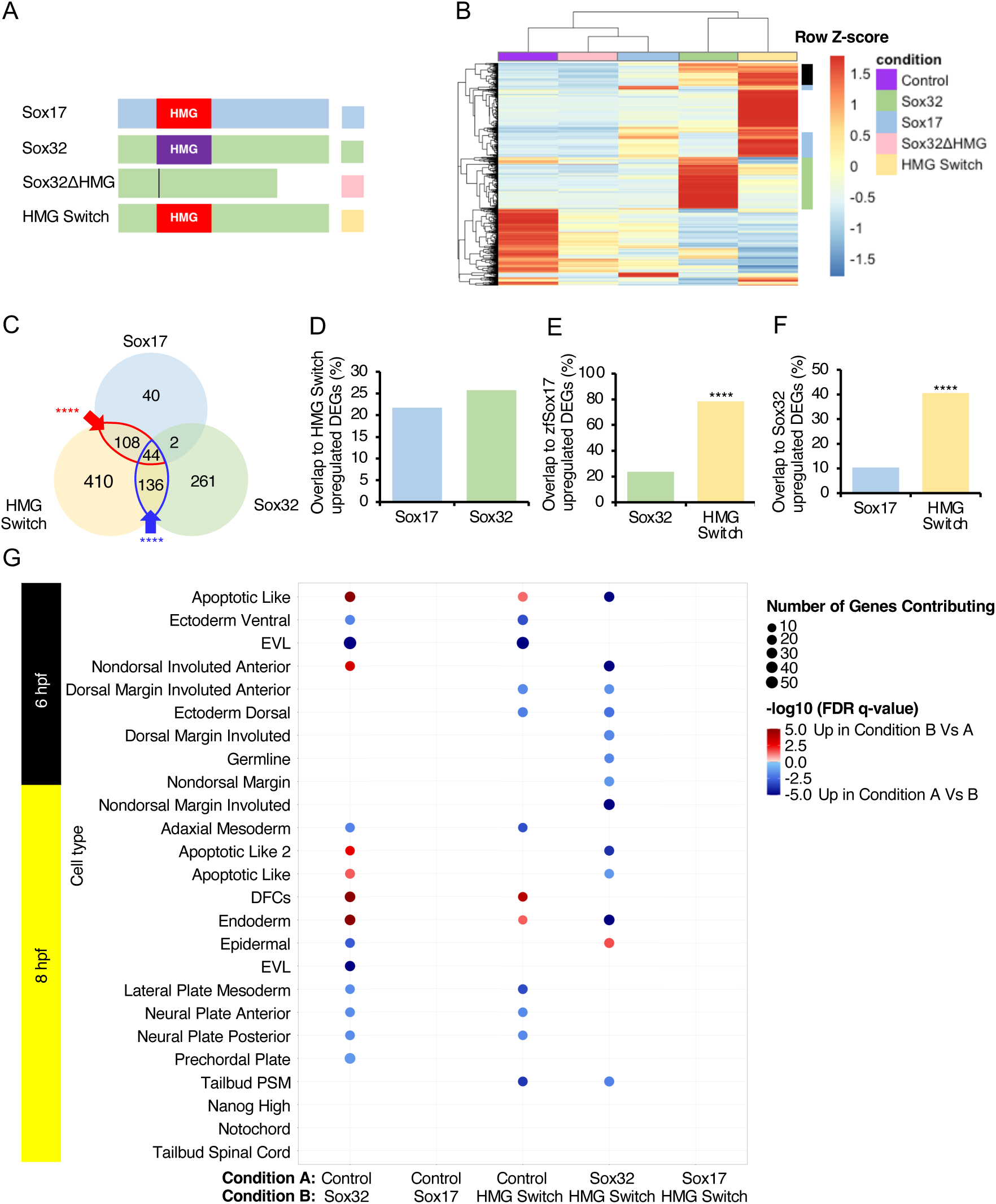
Differences between the Sox32 and Sox17 HMG domains are critical to the ability of Sox32 to induce endoderm and DFC gene expression. **A)** Protein models depicting constructs injected at the one-cell stage followed by RNA-sequencing at 6 hpf. **B)** Heatmap of all 1572 DEGs (Supplementary Tables S6-10), depicted as median counts from N=2 per condition. Green sidebar: induction of genes requiring the Sox32 HMG domain. Blue sidebar: induction of genes requiring the Sox17 HMG domain. Black sidebar: genes induced by Sox32 with either Sox32 or Sox17 HMG domain. **C)** Overlap of upregulated DEGs. Overlaps greater than expected according to Fisher’s Exact test are indicated - **** P < 0.0001. **D-F)** Percentage overlap of upregulated DEGs to (D) HMG Switch, (E) Sox17, (F) Sox32 upregulated DEGs. Bars represent the percentage of genes upregulated by the factor on the y-axis also upregulated by the factor on the x-axis. Statistical tests to determine whether the overlap was significantly greater with one factor on the x-axis compared to the other were carried out using Fisher’s Exact test.. **** P < 0.001. **G)** Gene Set Enrichment Analysis (GSEA) carried out against cell type markers defined by scRNA-seq at 6 and 8 hpf (Wagner, et al. 2018). EVL - enveloping layer; DFCs – dorsal forerunner cells; PSM – presomitic mesoderm.

As expected in the absence of a DNA-binding domain, Sox32ΔHMG failed to induce Sox32 target genes (Figure 3B). In contrast, HMG Switch-induced genes show significant overlap with both Sox32- and Sox17-induced genes, with either paralogue’s HMG domain sufficient for the induction of a core set of 180 target genes (Figure 3C). This is further supported in pairwise comparisons (Figure 3D-F). In fact, the majority of Sox17-induced genes are also induced by the HMG Switch protein (Figure 3C: 152 of 194 DEGs), underscoring the importance of the HMG domain for specific induction of Sox17 targets. However, the HMG Switch protein has a distinct TF profile, with over half (58.8%) of its target genes not induced by either Sox32 or Sox17. Furthermore, 58.9% of Sox32-induced genes are not induced by HMG Switch (Figure 3C). Taken together, this indicates that amino acid differences distinguishing the Sox32 and Sox17 HMG domains confer significant functional differences.

Functional annotation of induced target genes further illustrates the limits of the HMG Switch protein to recapitulate Sox32’s induction of specific cell identities (Figure 3G). Sox32 has a stronger capacity to significantly induce markers of endoderm and DFC cell types than the HMG Switch, and this enrichment occurs at the expense of other cell types such as the adaxial mesoderm and ventral ectoderm. This suggests the Sox32 HMG domain shows endodermal target specificity and is required to induce cell fate changes. Furthermore, while HMG Switch provides significant induction of endoderm markers compared to the control, Sox32 exhibits significantly greater induction of endoderm markers compared to HMG Switch (Figure 3G).

Overall, we conclude that the divergence of the Sox32 HMG domain from ancestral Sox17 is critical in conferring endoderm and DFC target specificity within the zebrafish embryo.

### The Sox32 25-aa C-terminal helix is required for induction of endoderm and DFC marker genes while the NTD has target-specific functions

Beyond the HMG domain, we also find evidence that the flanking protein regions influence target gene induction. One-fifth of Sox17’s targets are unique (Figure 3C), implicating other domains in determining the full repertoire of TF-specific target genes. This is further corroborated by the sizable overlap in induced targets between the HMG Switch and Sox32 proteins despite differing HMG domains (Figure 3C: 136 targets). We therefore next explored which peptides outside the Sox32 HMG domain are required for target gene induction.

The Sox32 N-terminal domain (NTD) contains a conserved EKR phosphorylation motif that, when phosphorylated in response to FGF signalling, leads to attenuation of Sox32 function, antagonising endoderm specification (Poulain 2006). Furthermore, examination of conservation across *sox32* orthologues revealed conservation of a 30-aa region within the Sox32 CTD, which we consider in two parts. Specifically, previous studies have identified a 7-aa β-catenin interacting peptide, with higher conservation between Sox32 and hSOX17 than Sox17 (Kikuchi, et al. 2001; Sinner, et al. 2004), that is required for Sox32 induction of ectopic endoderm upon OE (Zhao, et al. 2013). An upstream 23-aa peptide conserved among *sox32* orthologues remains unstudied (Supplementary Figure S5). Furthermore, AlphaFold predicts a high confidence 25-aa long CTD helix which partly encompasses these two peptide regions (Figure 2).

To examine the importance of these identified peptides to Sox32 target induction, we carried out specific domain deletions. We then analysed the ability of these domain-deleted mRNAs injected at the one-cell stage to induce five diagnostic endoderm and DFC cell marker genes relative to full length (FL)-Sox32 by 6 hpf (Figure 4A). All selected markers show induction by FL-Sox32 in RNA-seq experiments (Figure 1B), with tissue-specific expression in the endoderm only (*foxa2*) (Alexander, et al. 1999), endoderm and DFCs (*sox17, cxcr4a*) (Kikuchi, et al. 2000; Thisse 2001) or DFCs only (*vgll4l, ndr1*) (Rebagliati, et al. 1998; Thisse 2001).

**Figure 4:**
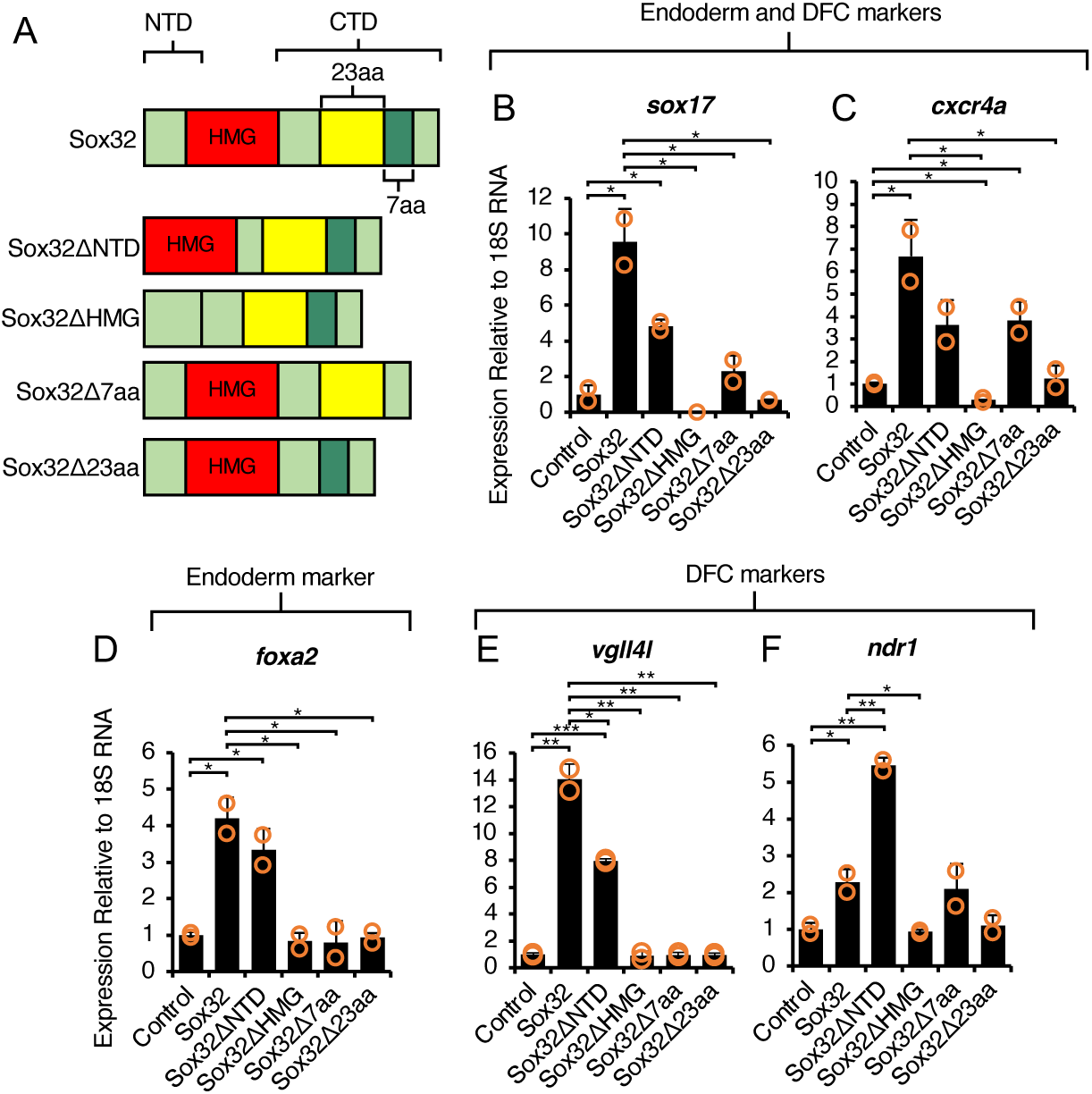
Putative helical CTD peptides highly conserved between Sox32 orthologues are required for induction of downstream endoderm and DFC marker genes, while NTD function is target-specific. **A)** Protein models of Sox32 deletions injected at the one-cell stage and RNA extraction followed by qPCR at 6 hpf. **B-F)** qPCR analysis of endoderm and DFC (B,C) endoderm only (D), and DFC only (E-F) markers. Bar charts depict the mean fold change relative to uninjected controls, error bars ± one standard deviation, orange circles indicate individual datapoints. Statistical tests on biological duplicate samples were carried out using Student’s *t*-test. * P<0.05, ** P <0.01, *** P<0.001.

Consistent with our global RNA-seq profiling (Figure 3B), Sox32 can strongly induce all marker genes and the HMG domain is essential (Figure 4B-F: FL-Sox32 and Sox32ΔHMG constructs). Equally, both peptides within the CTD helix are required, as Sox32Δ7aa and Sox32Δ23aa OE show no deviation in expression of target genes compared to control embryos (Figure 4B-F), except for significant induction of *cxcr4a* still attained with the Sox32Δ7aa construct.

The role of the NTD appears more complex, with differing roles depending on the target gene. Overall, the NTD would seem dispensable, as Sox32ΔNTD can significantly induce all analysed genes compared to the control, and with comparable levels of expression of the endoderm-only marker *foxa2* (Figure 4B). However, the Sox32ΔNTD construct only induced the dual endoderm/DFC markers *sox17* and *cxcr4a* to about half the level of FL-Sox32 (Figure 4C-D). Furthermore, for DFC-specific marker genes, relative induction by FL-Sox32 and Sox32ΔNTD differed, with significantly greater induction of *vgll4l* with the full-length protein but, conversely, significantly higher *ndr1* levels with the NTD-deleted construct (Figure 4E-F). Thus, the NTD may offer a tuneable interface for context-specific target gene regulation by Sox32, with the DFCs potentially evincing subtle modulation for different marker genes.

### Divergence of C-terminal peptides confers Sox32 and Sox17 differential target specificity

Sox32 requires its CTD helix for proper induction of its endodermal and DFC targets, specifically the 7-aa peptide (Figure 4), which is also necessary to induce ectopic endoderm upon Sox32 OE (Zhao, et al. 2013). This 7-aa region shows complete conservation to the zebrafish SoxF subfamily members Sox7 and Sox18 (Supplementary Figure S5) and high conservation to hSOX17. However, the 7-aa peptide is divergent in Sox17 (Figure 5A). This poses a question of whether this divergence between Sox32 and Sox17 confers differential target specificity. To decipher this, we generated constructs in which the respective 7-aa CTD regions were deleted from both Sox32 and Sox17, and hybrid constructs were also generated with 7-aa peptides switched between the two TFs (Figure 5B). Lastly, as the overall C-terminal helix structure also differs between Sox32 and Sox17 (Figure 2), we also produced a construct with the longer 25-aa Sox32 CTD helical peptide replacing the Sox17 7-aa peptide (Figure 5B).

**Figure 5:**
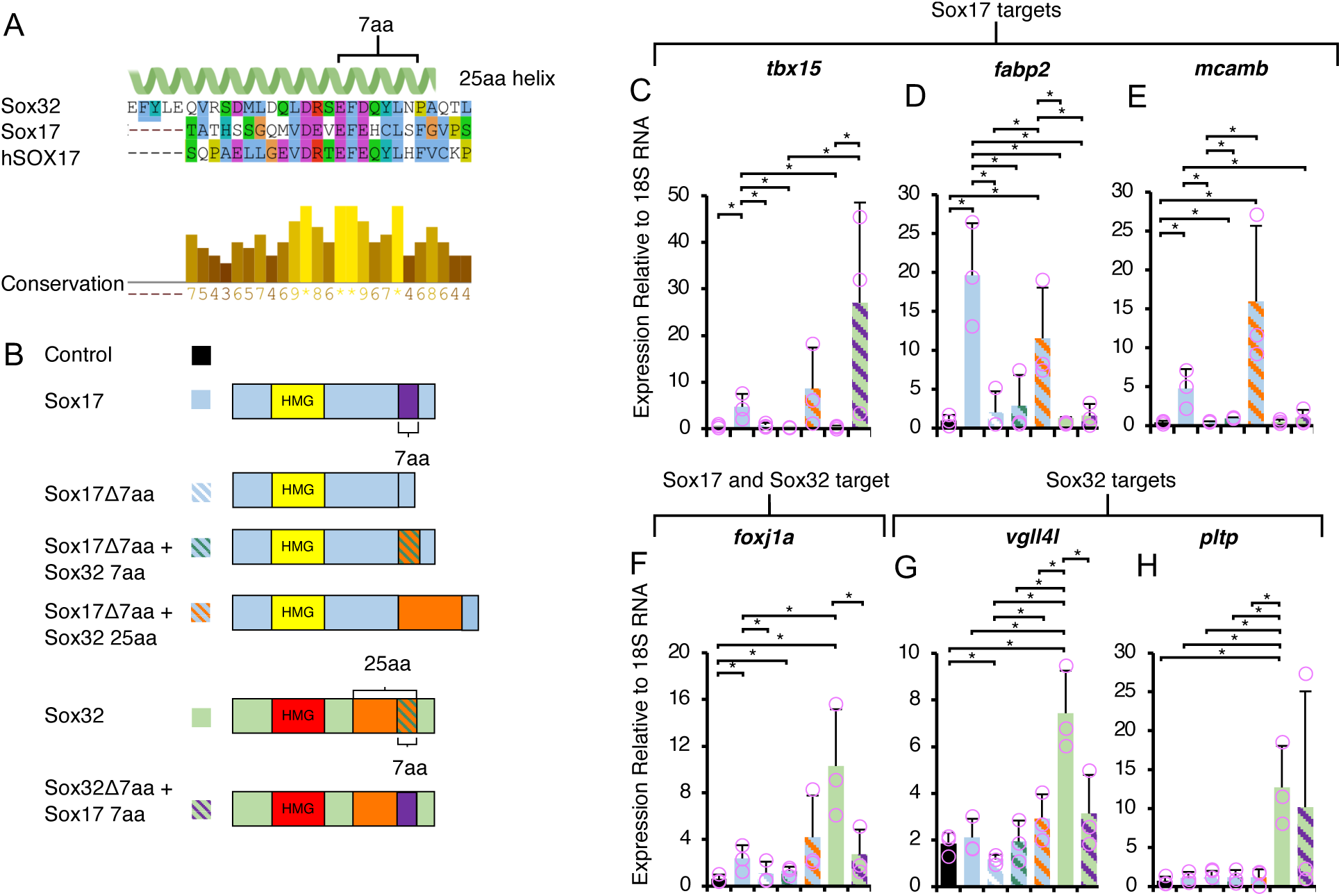
Divergence of 7-aa CTD putative helical peptides confers differential target specificity of Sox32 and Sox17. **A)** Alignment of Sox32 CTD helix (25 aa) to Sox17 and hSOX17 with conservation indicated. Alignment carried out using Clustal Omega in Jalview and colour coded based on conserved amino acid properties. Conservation levels calculated according to Analysis of Multiply Aligned Sequences (AMAS) displayed. **B)** Protein models of Sox17 and Sox32 domain deletions and switches injected at the one-cell stage and RNA extraction followed by qPCR at 6 hpf. **C-H)** qPCR of Sox17 targets (C-E), Sox17 and Sox32 targets (F) and Sox32 targets (G-H); sample condition colour code and order of bars corresponds to the legend in panel B. Bar charts depict the mean fold change relative to uninjected controls, error bars ± one standard deviation, pink circles indicate individual datapoints. Statistical tests on biological triplicate samples were carried out using Student’s t-test. * P<0.05.

Based on injections of each of the resulting mRNAs at the one-cell stage and qPCR analysis of target genes that distinguish Sox32 and Sox17 (Figure 1B), we find that induction of Sox17 targets indeed requires the Sox17 7-aa peptide. Replacement with the Sox32 7-aa fails to elicit FL-Sox17 levels of induction (Figure 5C-E). The replacement of the Sox17 7-aa peptide with the Sox32 25-aa peptide does not allow Sox17 to induce Sox32 targets (Figure 5G-H). Therefore, while the Sox32 CTD helix is necessary for Sox32 induction of endoderm and DFC targets (Figure 4), it is not sufficient in the context of the Sox17 protein. However, replacement of the Sox17 7-aa peptide with the Sox32 25-aa peptide does result in significant induction of Sox17 targets *fabp2* and *mcamb* compared to the control (Figure 5D-E). We therefore conclude that the Sox32 25-aa peptide that makes up the Sox32 CTD helix appears to be a potent transactivation domain.

In addition to replacement of the Sox17 7-aa peptide with that of Sox32 not inducing Sox17 targets to the capacity of FL-Sox17 (Figure 5C-E), we also see that replacement of the Sox32 7-aa peptide with the Sox17 7-aa peptide cannot induce *foxj1a* or *vgll4l* to the same level as FL-Sox32 (Figure 5G). This indicates that both Sox32 and Sox17 require their own intrinsic 7-aa domain for efficient induction of their respective targets. Therefore, we suggest that divergence in C-terminal peptides during evolution led to differential Sox32 and Sox17 target specificity and induction.

### Sox17 is necessary for establishing correct brain, heart and pancreas laterality

Given the distinct profile of Sox17 as a TF that is not merely redundant to Sox32, what is the full biological scope of its function? Organ development requires not only endoderm specification, but also subsequent tissue placement. For many species’ organ systems, bilateral asymmetry is vital for correct function, including in humans (Sutherland and Ware 2009; Agarwal, et al. 2021). Heterotaxia is a syndrome of abnormal abdominal and thoracic organ placement across the left-right axis of the body, including the pancreas and heart (Aiello, et al. 2007). In the brain, similar defects can alter patterns of gene expression and affect neural connectivity (Bianco and Wilson 2008).

Zebrafish Sox17 has previously been implicated in the establishment of LR asymmetry. Knockdown of Sox17 using an antisense morpholino (MO) was previously reported to result in abnormal pancreas placement, which could be rescued by co-injection with *sox17* mRNA to which the morpholino did not bind (Aamar and Dawid 2010). However, an analysis of zebrafish *sox17* function in genetic mutants is lacking. Furthermore, we aim to identify key domains of Sox17 required for correct LR patterning.

To further explore the role of Sox17 in LR asymmetry we used an efficient method of G0 CRISPR (Wu, et al. 2018) (Supplementary Figure S6) to disrupt the *sox17* locus by injection of Cas9-sgRNA complexes at the one-cell stage. We then analysed LR patterning using whole mount *in situ* hybridisation (WISH) of marker genes. We assessed organ placement for the normally dextrally looped heart (myocardial marker *myl7*, (Bakkers 2011) (Yelon, et al. 1999)), dextrally positioned pancreas (*insulin* expression, (Argenton, et al. 1999)), and sinistrally stronger gene expression in the habenular nuclei of the brain (*kctd12.1* expression, (Concha and Wilson 2001) (Sutherland 1982) (Gamse, et al. 2003; Gamse, et al. 2005)) (Figure 6A).

**Figure 6:**
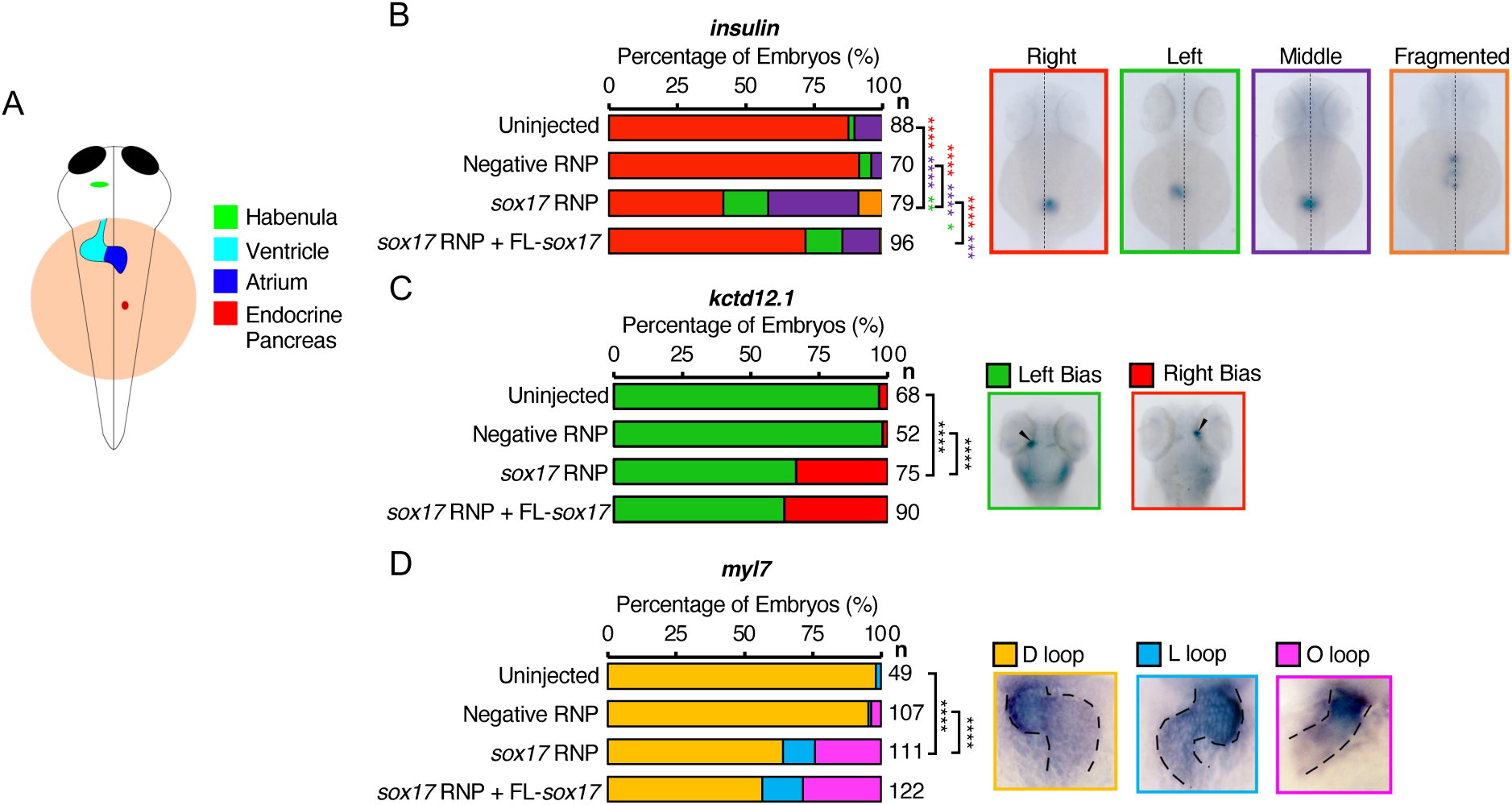
Sox17 CRISPants exhibit left-right asymmetry defects. **A)** Diagram depicting correct asymmetry of organs (4 days post-fertilisation - dpf). *Ktcd12.1* expression in the brain shows a left habenulae bias (green). The heart loops in a dextral (D-loop) fashion (blue). The endocrine pancreas is located to the right of the midline (red). **B-D)** WISH analysing the effect of Sox17 knockdown on endocrine pancreas placement relative to the midline (dotted line) at 48 hpf indicated by *insulin* (B); dorsal habenula *kctd12.1* asymmetric expression at 4 dpf – arrowheads indicate highest habenula expression (C); and heart looping at 48 hpf indicated by *myl7* – expression domain is outlined to highlight looping (D). Also shown are the phenotypic outcomes from rescue experiments from the co-injection of *sox17* exogenous mRNA along with CRISPR RNPs at the one-cell stage. Statistically significant differences in categorical scoring were inferred using Fisher’s Exact test on independent biological triplicate datasets. * P < 0.05, ** P < 0.01, *** P < 0.001, **** P < 0.0001. For B) red asterisks indicate a significant difference in right-sided pancreas *vs*. all other phenotypic outcomes, purple asterisks indicate a significant difference between middle-placed pancreas *vs*. everything else and green asterisks indicate a significant difference between left-sided pancreas and everything else. For C-D) black asterisks indicate significant difference between normal laterality and all other phenotypes. n= the number of embryos analysed.

We find that Sox17 CRISPants show abnormal brain asymmetry, heart looping and pancreas placement (Figure 6B-D). This includes a significant increase in the proportion of embryos exhibiting a middle and left-sided pancreas compared to the control (Figure 6B. We co-injected *sox17* mRNA into CRISPants at the one-cell stage to assess its ability to rescue the observed phenotypes. This resulted in a significant increase in embryos with normal right-sided pancreas placement. This occurred concurrently with a significant decrease in middle placement, while aberrant left-sided placement remained unaffected (Figure 6B). This indicates that the middle placement of the pancreas can be rescued by exogenous *sox17* mRNA, however left-sided pancreas, *i.e*., total reversal, cannot.

Neither brain asymmetry (Figure 6C) nor heart looping (Figure 6D) could be rescued by co-injection of exogenous *sox17* mRNA into CRISPants. To test whether exogenous Sox17 itself may be affecting LR patterning, we carried out OE studies through the injection of *sox17* mRNA across a broad range of concentrations. This indicated that Sox17 OE does not alter either brain asymmetry or heart looping (Supplementary Figure S7). The lack of rescue of abnormal phenotypes is therefore not due to the effect of Sox17 OE.

Going further, we considered whether the failure of *sox17* mRNA injection to rescue heart looping was due to underlying defects in adjacent tissue patterning. Sox32 mutants that lack anterior endoderm exhibit *cardia bifida* (Alexander, et al. 1999), a condition in which independent left and right heart fields develop due to a failure of coalescing morphogenesis. We therefore analysed our Sox17 CRISPants for expression of the anterior endoderm markers *foxa2* and *vgll4l.* Contrary to the reported Sox17 MO phenotype (Aamar and Dawid 2010), we found that the anterior endoderm is intact in Sox17 CRISPants (Supplementary Figure S8). Therefore, the lack of rescue of heart looping does not appear to be a secondary effect due to endoderm defects.

The failure of exogenous *sox17* mRNA to rescue defective pancreas positioning, brain asymmetry or heart looping led us to test whether KV function can be rescued. A key early step in the establishment of LR asymmetry, which is indicative of KV function, is unilateral *spaw* expression in the left LPM. Consistent with a defects in organ placement, we found that exogenous *sox17* mRNA injection at the one-cell stage failed to restore left-sided *spaw* expression in Sox17 CRISPants at 18 hpf (Supplementary Figure S9). This suggests that KV function is not restored by exogenous *sox17* mRNA. Since exogenous *sox17* mRNA is sufficient to restore right-sided placement of a proportion of pancreases that are aberrantly placed at the midline in Sox17 CRISPants, this suggests reduction of aberrant middle pancreas placement is independent of KV.

In conclusion, we have shown that Sox17 CRISPants have abnormalities in brain asymmetry, heart looping and pancreas placement. Only abnormal middle placed pancreas can be rescued by co-injection of the *sox17* mRNA into CRISPants. The placement of the pancreas is determined by looping of the posterior foregut as a result of LPM migration, in which *sox17* is also expressed (Horne-Badovinac, et al. 2003; Chung, et al. 2011). Therefore, Sox17 likely has a role in gut looping that exogenous *sox17* mRNA is sufficient to rescue, which we speculate may be linked to a function in the LPM.

### Conserved C-terminal peptides are required in Sox17 for correct pancreas placement

We have confirmed that Sox17 is required for organ asymmetry in zebrafish and that we can rescue pancreas placement through co-injection of *sox17* mRNA into CRISPants. We can therefore use this system to molecularly dissect which domains of Sox17 are required for its role in this process. We created *Sox17* mutant constructs with selected deletions based on high-level conservation among teleost and mammalian Sox17 orthologues (Supplementary Figure S10). Four conserved regions were identified and deleted: the Sox17 NTD, and three distinct regions within the CTD of varying lengths (Figure 7A: 35 aa, 14 aa and 20 aa). The mRNA corresponding to each mutant construct was injected into *sox17* CRISPants at the one-cell stage and their capacity to rescue correct pancreas placement was assessed.

**Figure 7:**
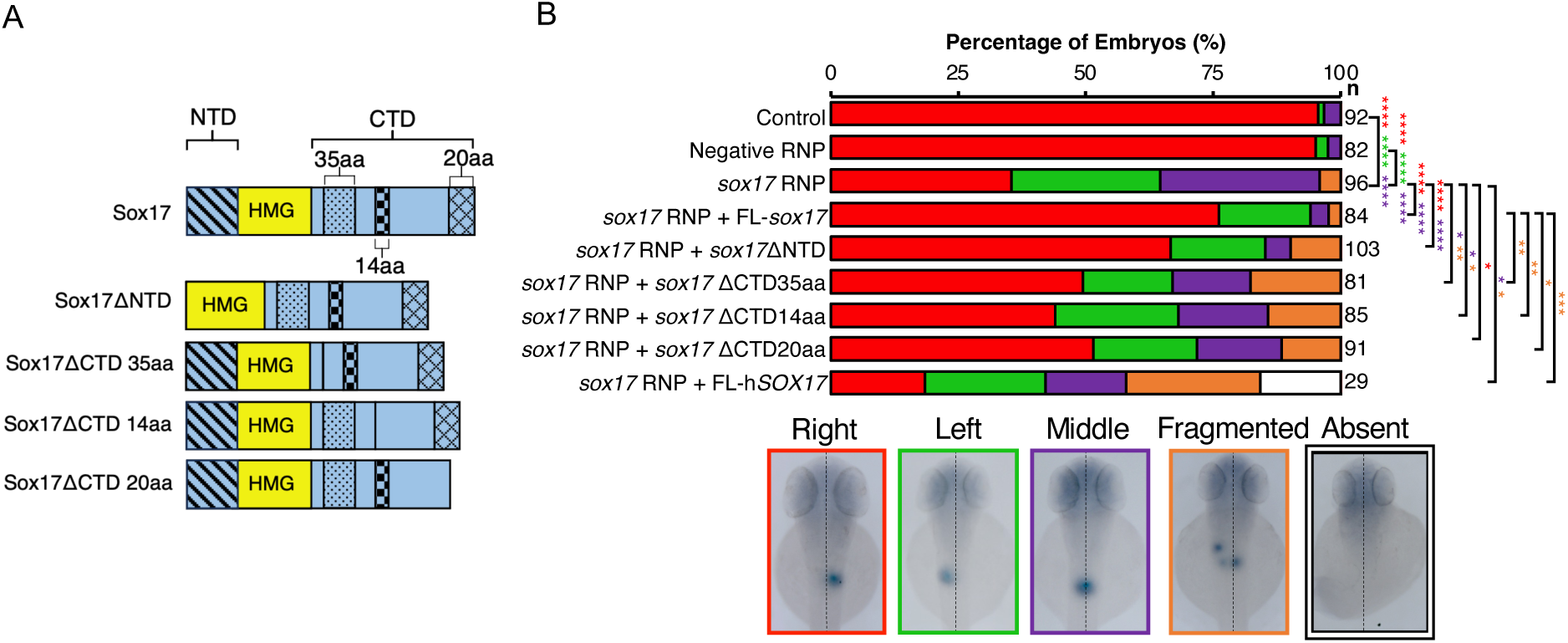
Sox17 CTD peptides are required for rescue of correct pancreas placement in Sox17 CRISPants. **A)** Diagram depicting protein models of Sox17 mutants. **B)** WISH at 48 hpf using *insulin* to assess the ability of Sox17 mutants in rescuing abnormal exocrine pancreas placement by co-injection of *sox17* exogenous mRNA alongside CRISPR RNPs at the one-cell stage. Statistically significant differences in categorical scoring were inferred using Fisher’s Exact test on independent biological triplicate datasets. * = P<0.05, ** = P<0.01, *** = P<0.001, **** = P<0.0001. Red asterisks indicate a significant difference in right-sided pancreas *vs.* all other phenotypes, purple asterisks indicate a significant difference between middle-placed pancreas *vs.* all other phenotypes, green asterisks indicate a significant difference between left-sided pancreas *vs.* all other phenotypes and yellow asterisks indicate a significant difference between fragmented *insulin*+ domains *vs.* all other phenotypes. FL refers to full length. n= the number of embryos analysed.

We find that Sox17ΔNTD can rescue pancreas placement in *sox17* CRISPants to a similar degree as full length (FL)-Sox17. Both constructs result in a significant increase in CRISPants exhibiting normal right-sided pancreas placement and a significant decrease in embryos showing middle placement, with left-sided placement unaffected compared to *sox17* CRISPants alone (Figure 7B). This indicates the N-terminal domain of Sox17 is largely dispensable for its function in pancreas placement. Sox17ΔCTD35aa and Sox17ΔCTD14aa mRNA injections lead to a significant decrease in *sox17* CRISPant embryos with a middle-placed pancreas. However, this does not coincide with an increase in normal right-sided placement as seen with FL-Sox17 and Sox17ΔNTD. Instead, there is a significant increase in embryos exhibiting multiple fragmented *insulin*+ domains (Figure 7B). Sox17ΔCTD20aa mRNA injection leads to a significant increase in CRISPants with normal right-sided placement compared to embryos injected with *sox17* RNP alone, suggesting this mutant construct retains some degree of relevant function. However, we also note that we reduction in aberrant pancreas placement is also accompanied by a moderate increase in pancreas fragmentation (Figure 7B). Overall, all three CTD mutants may have some reduced capability to restore migration of the pancreas to the right, though not to the degree of FL-Sox17 and Sox17ΔNTD. However, the CTD mutants also disrupt pancreas formation, either by driving ectopic specification of pancreas identity, or migration defects leading to the fragmented phenotype observed.

*Sox17*^-/-^ mutant mice exhibit an open body wall and failure to turn from a concave to a convex “foetal” position (Kanai-Azuma, et al. 2002; Viotti, et al. 2012) – characteristics of mutants with LR patterning defects (Hamada, et al. 2002). Given the requirement for Sox17 in LR patterning in mammals and zebrafish, we sought to determine whether mammalian SOX17 has the capacity to induce correct pancreas placement through injection of h*SOX17* mRNA into *sox17* CRISPants. We found that hSOX17 is unable to rescue pancreas placement. Instead, it leads to a further decrease in right-sided pancreas placement in CRISPants, or even the total absence of the pancreas (Figure 7B). The aberrant LR phenotype in mouse has been suggested to be largely a secondary effect due to depletion of gut endoderm (Viotti, et al. 2012), while zebrafish Sox17 CRISPants do not exhibit loss of endoderm (Supplementary Figure S7). Taken together, this indicates that while Sox17 is required for LR asymmetry in mammals and zebrafish, mechanistic variability exists.

Overall, we conclude the Sox17 CTD is required for correct pancreas placement, while the NTD is dispensable. Additionally, our results indicate that peptides in the CTD can influence formation of an intact endocrine pancreas, although whether this reflects aberrant morphogenesis or specification is unclear.

Taken together, our results reveal the molecular basis for functional divergence between Sox17 and Sox32, highlighting key functional peptides controlling both endoderm and DFC specification, and the establishment of pancreas placement during gut morphogenesis.

## Discussion

A classical view of the fate of duplicated genes that remain functional during evolution is that they either undergo sub- or neofunctionalization (Tasnim, et al. 2024). How does the case of Sox17 and Sox32 reflect this? The expression domains of *sox32* and *sox17* substantially overlap in the early embryo with Sox32 activating *sox17* expression, and are similar to *Sox17* expression patterns in other species. We therefore consider sub- or neofunctionalization in the context of protein function rather than expression domains.

### Divergence of the Sox32 HMG domain is critical to allow engagement with the teleost endoderm GRN

Consistent with previous reports (Kikuchi, et al. 2001; Stafford, et al. 2006) our analyses indicate that ectopic Sox32 expression induces endodermal cell identity at the expense of other cell types, indicating its ability to change cell fate (Figure 3G). Remarkably, it also induces a significant number of DFC marker genes suggesting that while Sox32 is necessary for DFC identity it is also substantially sufficient to direct the DFC gene expression programme (Figure 3G).

Our molecular analyses reveal that the ability of Sox32 to orchestrate the endoderm GRN is dependent on differences between the Sox32 and Sox17 HMG domains. Given the greater similarity between Sox17 and hSOX17 HMG domains, we suggest Sox32 diverged after the duplication event where *sox32* and *sox17* emerged. This suggests a scenario wherein subfunctionalisation has occurred with Sox32 adopting the key role in endoderm formation, while Sox17 retains other roles such as regulating genes involved in vasculogenesis, as SOX17 does in mammals (Figures S2, S4). However, the divergence of the Sox32 HMG domain and the inability of hSOX17 to induce endoderm marker expression in zebrafish raises questions about the mechanistic similarities by which these TFs orchestrate endoderm gene expression in their native species. It is possible that the different HMG domains bind different *cis*-regulatory modules (CRMs) and hence activate different downstream target genes in our experiments, either due to different DNA binding preferences or altered physiochemical interactions with co-factors. However, all Sox TFs are considered to individually bind similar sequence motifs (Kondoh and Kamachi 2010). Furthermore, contact sites between mouse SOX17 HMG domain and a *cis*-regulatory enhancer upstream of the *Lama1* transcriptional start site have been mapped (Palasingam, et al. 2009) and are highly conserved in Sox32 and Sox17 (Figure 2D). This may indicate that Sox32, Sox17 and hSOX17 HMG domains are individually capable of engaging similar DNA sequences. A more likely scenario is that the differences between HMG domains alters the spectrum or configuration of co-factor interactions, impacting co-operative engagement of TF complexes with CRMs. However, it does not necessarily follow that hSOX17 and Sox32 have substantially different co-factors during endoderm formation if orthologous co-factors have undergone parallel evolution. In this scenario hSOX17 and Sox32 would only be able to interact with co-factors from their native species, which would preclude hSOX17 from inducing endoderm marker expression in zebrafish. It would also explain why the Sox17 HMG domain cannot substantially engage the endoderm GRN, if key co-factors have evolved to engage Sox32.

While very few Sox32 interacting TFs have been identified, an interesting candidate that may explain the differences in HMG domain functions is the TF Pou5f3. Pou5f3 is required for Sox32-mediated induction of endoderm fate (Lunde, et al. 2004; Reim, et al. 2004; Zhao, et al. 2013). However, in mammals the SOX17 HMG domain interacts with the POU domain of POU5F1 to cooperatively bind CRMs and orchestrate target gene induction (Stefanovic, et al. 2009; Ng, et al. 2012). While not true orthologues, human *POU5F1* is the closest homologue of zebrafish *pou5f3*. Duplication of an ancestral *Pou5* gene occurred in the last common ancestor of extant cartilaginous fishes and bony fishes leading to two genes. Subsequent gene loss has led to maintenance of only *pou5f3* in teleosts and *POU5F1* in eutherians including humans (Frankenberg and Renfree 2013). POU5F1 (also known as OCT3 or OCT4) is known to interact with the HMG domains of many SOX factors (Ng, et al. 2012). However, distinctions between HMG domains leads to alterations in POU5F1 engagement, and consequently different SOX-POU5F1 target sequence selection (Ng, et al. 2012). Thus, altered configuration of Sox32-Pou5f3 interactions compared to SOX17-POU5F1 would likely prevent engagement with the same CRMs. Similarly, the requirement for correct partner cooperation in target transcription has been established in human SOX TF function. For example, SOX2 can interact with both OCT3 and OCT1, but only the SOX2-OCT3 complex can promote transcriptional activation of the FGF4 enhancer (Yuan, et al. 1995). It is therefore likely that Sox32 complexes have different sequence specificities to Sox17- or hSOX17-containing complexes, either through altered interaction with the same/homologous co-factors or altered co-factor selection. This would lead to different target site selection and distinct functional outcomes for SOX factors that are otherwise capable of binding the same individual consensus sequence. Future analyses should therefore focus on the interacting partners and DNA binding preferences of resulting Sox TF complexes, and whether hSOX17 and Sox32 have demonstrably similar or distinct target CRMs in their native species.

### Functional partition and divergence through alteration of C-terminal helices

A key distinction between these endoderm-relevant SoxF factors is a putative CTD helix, which is notably longer in Sox32 than Sox17 (Figure 2A-C). The 25-aa high confidence prediction in Sox32 (Figure 2A) encompasses a short peptide (EF<u>D</u>QYL) which deviates from hSOX17 by only a single amino acid of similar properties (EF<u>E</u>QYL). Importantly, this EF(D/E)QYL motif was first identified in *Xenopus* where Sox17α shows 100% sequence identity to hSOX17 and Sox17β to Sox32, with both peptides critical to Sox17αβ interaction with Wnt pathway effector β-catenin (Sinner, et al. 2004). Additionally, *Xenopus* Sox17α/β and hSOX17 co-occupy enhancers with β-catenin resulting in activation of endoderm-specific genes and repression of mesoectodermal genes (Mukherjee, et al. 2020; Mukherjee, et al. 2022). However, zebrafish Sox17 exhibits alterations in this peptide including replacement of the central polar uncharged glutamine (Q) residue with a positively charged histidine (H) residue, and also substitution of the polar amino acid tyrosine (Y) for non-polar cysteine (C) (Figure 2E). It is therefore likely that Sox32 has retained the ancestral function of recruiting β-catenin to activate endoderm-specific genes, while divergence of this peptide in Sox17 has led to neofunctionalisation. This is supported by our observation that both Sox32 and Sox17 require their own individual peptide sequences (EFDQYLN and EFEHCLS respectfully as tested) for induction of their distinct target genes. Thus, the function of Sox32 in directing the endoderm GRN relies on divergence of its HMG domain and retention of the C-terminal β-catenin interacting peptide, while alterations to this C-terminal peptide in Sox17 likely led to altered target gene selection and consequent functional diversification.

The 25-aa CTD helical domain in Sox32 contains 18 aa upstream of the β-catenin interacting peptide. Our analyses (via deletion of a 23 aa peptide encompassing the 18-aa helix and five upstream amino acids) indicates that this broader helical domain is also essential for Sox32-mediated induction of endoderm and DFC target genes (Figure 4). Additionally, though Sox17 requires its divergent EFEHCLS peptide for induction of its target genes, substitution for the Sox32 25-aa CTD helical domain nevertheless leads to potent induction of Sox17 (though not Sox32) target genes (Figure 5D-E). We therefore hypothesise the Sox32 CTD helix is a potent transactivation domain. Interestingly, C-terminal helices in FoxA factors have been shown to be critical for gene activation, facilitating binding to histone octamers leading to nucleosome eviction and increased chromatin accessibility (Pani, et al. 1992; Iwafuchi, et al. 2020). It would be interesting to determine whether the 25 aa CTD helix in Sox32 similarly confers pioneer TF activity.

### Sox17 potentially has multiple roles in establishing organ asymmetry

Our analyses of G0 CRISPants suggests Sox17 is necessary for correct unilateral Nodal (*spaw*) expression, brain asymmetry, heart looping and pancreas placement (Figure 6). Furthermore, we have shown that co-injection with exogenous *sox17* mRNA is sufficient to rescue abnormal pancreas placement phenotypes in Sox17 CRISPants, consistent with previous experiments with antisense morpholinos (Aamar and Dawid 2010). However, exogenous *sox17* mRNA is not sufficient to rescue abnormal Nodal expression, heart looping or brain asymmetry. This tantalisingly suggests Sox17 may have distinct roles in different cell types to establish organ asymmetry, as we discuss below.

The process of zebrafish heart morphogenesis and looping has been extensively studied and reviewed (Bakkers 2011). While we have shown that an intact *sox17* locus is necessary for correct zebrafish heart looping, the lack of phenotypic rescue by exogenous *sox17* mRNA cannot be attributed to injection of exogenous *sox17* mRNA (Supplementary Figure S7), or a secondary consequence of an endodermal phenotype (Supplementary Figure S8). Our RNA-seq analyses show Sox17-induced genes are enriched for heart-related anatomical terms, including endocardium and ventricular myocardium (Supplementary Figure S2). Additionally, we have observed *sox17:EGFP* reporter expression in the heart including in the endocardium, based on co-expression with *kdrl:mCherry* (Supplementary Figure S4A, Video S1). These cells arise from precursors in the LPM, which is *sox17*+ (Zhong, et al. 2001; Chung, et al. 2011). *Sox17* is expressed in mouse endocardium, where it is required for proper heart morphogenesis (Saba, et al. 2019). Considering we find *sox17:EGFP*+ cells within the zebrafish heart (Supplementary Figure S4), a conserved role within the endocardium controlling heart morphogenesis is plausible.

Sox17 is also expressed within KV (Schneider, et al. 2008; Chung, et al. 2011), and morpholino knockdown of Sox17 leads to abnormality in KV morphogenesis and function via defective ciliogenesis (Aamar and Dawid 2010). The perturbation of cilia function results in aberrant *spaw* expression within the LPM (Kramer-Zucker, et al. 2005). We show that Sox17 CRISPants exhibit aberrant *spaw* expression in the LPM, and this cannot be restored to normal left-sided *spaw* expression through co-injection of exogenous *sox17* mRNA (Supplementary Figure S9). This suggests we are not able to rescue KV function with exogenous *sox17* mRNA, leading to an inability to rescue correct brain asymmetry and heart looping. An explanation may be off-target disruption of another gene acting in KV function. However, this seems unlikely given morpholino knockdown of Sox17 also leads to KV defects (Aamar and Dawid 2010). Alternatively, since KV typically consists of ∼50 cells (Gokey, et al. 2016) – a tiny proportion of the embryo – it is possible that the failure to rescue left-sided *spaw* expression is due to a lack of inheritance of exogenous *sox17* mRNA by sufficient KV cells, as *sox17* OE itself does not lead to LR defects (Supplementary Figure S7).

However, the failure to rescue KV function (and thus brain asymmetry and heart looping) makes the rescue of pancreas placement even more tantalising. As well as KV function, visceral organ laterality and therefore pancreas placement is also determined by LPM migration, which pushes the developing pancreas to the right of the midline due to left-sided Nodal signalling (Horne-Badovinac, et al. 2003). Alterations in Nodal signalling within the LPM randomises the pattern of LPM migration, resulting in aberrant pancreas placement (Horne-Badovinac, et al. 2003). Remarkably, however, exogenous *sox17* mRNA can restore gut looping in Sox17 CRISPants as indicated by a significant reduction in midline-placed pancreas, even though it cannot rescue KV function (Figure 6B). Since *sox17* is expressed in LPM (Chung, et al. 2011), this suggests that LPM migration is compromised in Sox17 CRISPants, but can be restored through exogenous *sox17* mRNA injection. It is tempting to speculate that the fragmented pancreas phenotype we observe on mutant *sox17* mRNA injection also reflects effects on LPM migration (though ectopic pancreas specification is also a possibility). Overall, our data confirms Sox17 controls LR asymmetry through roles in KV, and suggests a new role for Sox17 in gut looping and possibly also in heart looping through expression in the endocardium. Our analyses show that multiple peptides conserved among Sox17 orthologues are required for rescue of the pancreas placement defect in Sox17 CRISPants. Strikingly, amongst these is the Sox17CTDΔ14 aa deletion which encompasses the EFEHCLS sequence noted to have diverged from the ancestral β-catenin interacting peptide. It is therefore possible that this divergence of Sox17 is key to its role in controlling gut looping in zebrafish.

Overall our results indicate that evolutionary alterations in the HMG domain of Sox32 combined with the presence of a CTD helical domain including retention of a β-catenin interacting peptide are key to its orchestration of the endoderm and DFC GRNs. While hSOX17 clearly cannot engage the zebrafish endoderm GRN in the context of these experiments, it would be interesting to explore whether Sox32 and hSOX17 have similar target genes and co-factors within their native species. In contrast to Sox32, alterations in the CTD of Sox17 have driven functional divergence. This alters target gene specificity in the early embryo and may be related to a new function for Sox17 in controlling gut looping to establish visceral organ asymmetry, which warrants further in-depth investigation.

## Materials and Methods

### Animals

AB, *Tg(sox17:EGFP)^ha01Tg^* and *Tg(kdrl:mCherry^S896^,gata1a:dsRed^sd2tg^)* fish were reared as described (Westerfield 2000; Traver, et al. 2003; Chi, et al. 2008; Mizoguchi, et al. 2008). All zebrafish studies were conducted in compliance with the United Kingdom Animals (Scientific Procedures) Act 1986, licensed by the UK Home Office and implemented by The University of Warwick.

### Confocal Microscopy

48 hpf zebrafish embryos were anesthetized with 0.168%(w/v) of Ethyl 3-aminobenzoate methanesulfonate, prior to mounting in 1% lateral view agarose moulds. Moulds were generated from a 3D stamp as designed by (Kleinhans and Lecaudey 2019). To ensure high-quality imaging, embryos were treated before 24 hpf with 0.003% (w/v) 1-Phenyl-2-thiourea (PTU) to prevent pigmentation (Karlsson, et al. 2001). Imaging was carried out on a Zeiss LSM 980 with Airyscan 2.

### Cloning for *in vitro* production of mRNAs

pCS2+*hSOX17* was cloned from PB-TRE3G-SOX17 (Miller, et al. 2018) using XhoI and XbaI restriction sites. PB-TRE3G-SOX17 was a gift from David Vereide (Addgene plasmid # 104541; http://n2t.net/addgene:104541; RRID:Addgene_104541) *Sox17* and *sox32* deletion and domain switch constructs (Supplementary Table S11) were generated using pCS2+*sox17* and pCS2+*myc-sox32* by PCR amplification using Q5 polymerase (NEB) with described primers (Supplementary Table S12) followed by re-circularisation or insertion using T4 ligase (Invitrogen). All constructs were generated using blunt-end cloning, except for Sox17Δ7aa + Sox32 25aa and Sox32ΔHMG + Sox17 HMG where NEBuilder HiFi DNA Assembly (NEB), with Sox17Δ7aa + Sox32 25aa as a gBlock (Integrated DNA Technologies) was used.

### mRNA *in vitro* transcription

Capped mRNA was synthesised from pCS2+ plasmids containing genes of interest. Linearisation with NotI was conducted followed by transcription using SP6 RNA polymerase as previously described (Talbot, et al. 2022).

### Ribonuclear protein (RNP) complex production

To knockdown *sox17* ribonucleoprotein (RNP) complexes were generated using a pool of three guide RNAs and Cas9 protein (Integrated DNA Technologies) as described (Essner 2016). Guide RNAs that do not target the zebrafish genome were used to generate a negative control RNP. Guide RNAs were generated from specific crRNAs (Supplementary Table S13) and standard tracrRNA (Integrated DNA Technologies) as described (Essner 2016). crRNAs against *sox17* were predesigned by Integrated DNA Technologies and selected based on predicted on-target and off-target scores from both Integrated DNA Technologies and CHOPCHOP (Labun, et al. 2019).

### Microinjection

Equimolar amounts of mRNA (calculated based on relative mRNA length) were injected at the one-cell stage. For interspecies RNA-seq each embryo was injected with the following amount: *sox32* – 150pg, h*SOX17* – 141pg, *sox17* – 175pg. For HMG Switch RNA-seq each embryo was injected with the following amount: *sox32* – 150pg, *sox32*ΔHMG – 121.5pg, and *sox32*ΔHMG+*sox17*HMG – 148.5pg. For *sox32* mutant qPCR, each embryo was injected with the following amount: *sox32* – 150pg, - *sox32*ΔNTD – 127.5pg, *sox32*ΔHMG – 121.5pg, *sox32*ΔCTD23aa – 142.5pg, *sox32*ΔCTD7aa – 147pg. For CTD switch qPCR, each embryo was injected with the following amount: *sox17* – 150pg, *sox17*Δ7aa – 147pg, *sox32* – 128ng, *sox32*Δ7aa+*sox17-*7aa – 128pg, *sox17*Δ7aa+*sox32-*7aa – 150pg, *sox17*Δ7aa+*sox32-* 25aa – 155pg. For Sox17 functional dissection, each embryo was injected with the following amount of mRNA: *sox17* – 150pg, *sox17*ΔNTD – 134pg, *sox17*ΔCTD35aa – 139pg, *sox17*ΔCTD14aa – 145pg, *sox17*ΔCTD20aa – 143pg, h*SOX17* – 108pg. For overexpression experiments mRNA was injected in a volume of 1nL. To produce CRISPant embryos RNPs were injected in a volume of 3nL. For experiments involving combined CRISPR RNP and mRNA injection RNPs and mRNAs were injected in a combined volume of 4nL.

### Whole mount *in situ* hybridisation (WISH)

WISH of zebrafish embryos was carried out as described (Thisse and Thisse 2008). Anti-sense RNA probes for *myl7* (Yelon, et al. 1999), *insulin* (Milewski, et al. 1998), *vgll4l* (Nelson, et al. 2014) and *foxa2* (Odenthal and Nüsslein-Volhard 1998) were synthesised as described. *Ktcd12.1* RNA probe was generated from a PCR product using primers: F - CTGCCGGACTATTTTCCAGAG, R-TAATACGACTCACTATAGGGAGCTGCACGCGACCATCT and transcribed using T7 polymerase (Promega). Categorical scoring on randomised samples was completed blinded.

### Total RNA Extraction

Embryos were lysed at 6 hpf via addition of 600ul buffer RLT and DTT (2% v/v) to at least fifty embryos per condition, with initial disruption using a plastic pestle. Further disruption was carried out using QIAshredder columns (QIAGEN) and RNA was extracted using the RNeasy Mini Kit (QIAGEN) with on-column DNase treatment (QIAGEN) as per manufacturer’s protocol.

### qPCR

cDNA was generated from 400ng of total RNA using Random Hexamers (Promega) and Ultrascript Reverse Transcriptase (PCR Bio) as per manufacturer’s protocol. qPCR was carried out using relevant primers (Supplementary Table S14) and Power SYBR Green Master Mix (Thermofisher) on the Stratagene Mx3005P (Agilent Technologies). Raw Ct values were extracted using MxPro software and analysed using the τιτιCt method (Livak and Schmittgen 2001), with normalisation to *18S* rRNA (McCurley and Callard 2008). Statistical differences were inferred using Students t-test with significance set at P <0.05.

### RNA-sequencing Library Preparation and Analysis

Total RNA was sent to the Cologne Center for Genomics (CCG) for library preparation (Illumina TruSeq Stranded mRNA) and sequencing on the Illumina NovaSeq 6000 platform with a read length of 101 bp.

Quality control of raw data was performed using FastQC. Reads were mapped to danRer11 zebrafish genome and counts per gene (Ensembl version 97) generated using STAR with default settings (Dobin, et al. 2013). Differential expression was identified through adjusted P-value < 0.05 using DESeq2 (1.36.0) (Love, et al. 2014) in R (4.2.0). To identify whether different TFs illicit similar or different effects on the transcriptome, upregulated and downregulated DEGs were separately overlapped to identify whether TFs induce or repress similar genes. Statistical testing to identify whether the overlap in genes was significantly more/less than expected by chance was conducted using Fisher’s exact test (phyper in R). This analysis used a gene universe size defined as all genes detected in RNA-seq where samples in all conditions showed counts > 0. Heatmaps were generated on median normalised counts per condition using pheatmap (1.0.12) with ‘scale = row’ option to display deviation in expression from the average expression of each gene. Gene Ontology (GO) and anatomy term enrichment analysis was carried out on fishEnrichR (Chen, et al. 2013; Kuleshov, et al. 2016) using autoRIF Z-score, with significantly enriched terms defined by an adjusted P-value < 0.05.

To infer how TFs influence cell fate, Gene Set Enrichment Analysis (GSEA) was carried out using published scRNA-sequencing data (Wagner, et al. 2018) as cell identity markers using GSEA (4.3.2) (Mootha, et al. 2003; Subramanian, et al. 2005).

### Conservation Analysis

Protein sequences were obtained from UniProt (The UniProt 2023) (Supplementary Table S15) and alignments were conducted aligned using Clustal Omega in JalView (Sievers, et al. 2011; Sievers and Higgins 2018). Jalview uses multiple sequence alignments according to Analysis of Multiply Aligned Sequences (AMAS) (Livingstone and Barton 1993) to calculate conservation levels. Conserved sequences were compared to 3D structures obtained from AlphaFold; Sox32 (Q90Z46), Sox17 (Q5PQZ5), hSOX17 (Q9H6I2) (Jumper, et al. 2021; Varadi, et al. 2022).

## Authors’ contributions

Conceived and designed the experiments: ACN, SJ, KAP. Performed the experiments: SJ, RE. Analysed the data: SJ. Writing – original draft preparation: SJ. Writing – review and editing: ACN, KAP.

## Acknowledgments

We thank Rui Monteiro for kindly gifting us the *Tg(kdrl:mCherry,gata1a:dsRed)* fish used in this study. We thank Fiona Wardle and Karuna Sampath for reagents. We thank Karuna Sampath, Emily Noël and Andre Pires da Silva for valuable discussions, and Karuna Sampath and Andre Pires da Silva for generous access to equipment. We thank Eirini Vlachaki for technical support early in the project. We also thank the Warwick zebrafish facility for zebrafish care and, Ian Hands-Portman from the Warwick School of Life Sciences for imaging advice and support.

## Data Accessibility

All RNA-seq data have been submitted to NCBI Gene Expression Omnibus under accession number GSE274063.

## References

Aamar E, Dawid IB. 2010. Sox17 and chordin are required for formation of Kupffer’s vesicle and left-right asymmetry determination in zebrafish. Developmental dynamics : an official publication of the American Association of Anatomists 239:2980–2988.

Agarwal R, Varghese R, Jesudian V, Moses J. 2021. The heterotaxy syndrome: associated congenital heart defects and management. Indian Journal of Thoracic and Cardiovascular Surgery 37:67–81.

Aiello VD, Anderson RH, Béland MJ, Colan SD, Duca DD, Elliott MJ, Franklin RCG, Jacobs JP, Krogmann ON, Kurosawa H, et al. 2007. The nomenclature, definition and classification of cardiac structures in the setting of heterotaxy. Cardiology in the Young 17:1–28.

Alexander J, Rothenberg M, Henry GL, Stainier DYR. 1999. casanova Plays an early and essential role in endoderm formation in zebrafish. Developmental Biology.

Angelozzi M, Lefebvre V. 2019. SOXopathies: Growing Family of Developmental Disorders Due to SOX Mutations. Trends Genet 35:658–671.

Aoki TO, David NB, Minchiotti G, Saint-Etienne L, Dickmeis T, Persico GM, Strähle U, Mourrain P, Rosa FdrM. 2002. Molecular integration of casanova in the Nodal signalling pathway controlling endoderm formation. Development 129:275–286.

Argenton F, Zecchin E, Bortolussi M. 1999. Early appearance of pancreatic hormone-expressing cells in the zebrafish embryo. Mechanisms of Development 87:217–221.

Bakkers J. 2011. Zebrafish as a model to study cardiac development and human cardiac disease. Cardiovascular Research 91:279–288.

Bianco IH, Wilson SW. 2008. The habenular nuclei: a conserved asymmetric relay station in the vertebrate brain. Philosophical Transactions of the Royal Society B: Biological Sciences 364:1005–1020.

Bowles J, Schepers G, Koopman P. 2000. Phylogeny of the SOX Family of Developmental Transcription Factors Based on Sequence and Structural Indicators. Developmental Biology 227:239–255.

Chen EY, Tan CM, Kou Y, Duan Q, Wang Z, Meirelles GV, Clark NR, Ma’ayan A. 2013. Enrichr: interactive and collaborative HTML5 gene list enrichment analysis tool. BMC Bioinformatics 14:128.

Chi NC, Shaw RM, De Val S, Kang G, Jan LY, Black BL, Stainier DYR. 2008. Foxn4 directly regulates tbx2b expression and atrioventricular canal formation. Genes & development 22:734–739.

Chung MIS, Ma ACH, Fung T-K, Leung AYH. 2011. Characterization of Sry-related HMG box group F genes in zebrafish hematopoiesis. Experimental Hematology 39:986–998.e985.

Concha ML, Wilson SW. 2001. Asymmetry in the epithalamus of vertebrates. Journal of Anatomy 199:63–84.

Cooper MS, D’Amico LA. 1996. A Cluster of Noninvoluting Endocytic Cells at the Margin of the Zebrafish Blastoderm Marks the Site of Embryonic Shield Formation. Developmental Biology 180:184–198.

Dickmeis T, Mourrain P, Saint-Etienne L, Fischer N, Aanstad P, Clark M, Strähle U, Rosa F. 2001. A crucial component of the endoderm formation pathway, CASANOVA, is encoded by a novel sox-related gene. Genes & development 15:1487–1492.

Dobin A, Davis CA, Schlesinger F, Drenkow J, Zaleski C, Jha S, Batut P, Chaisson M, Gingeras TR. 2013. STAR: ultrafast universal RNA-seq aligner. Bioinformatics (Oxford, England) 29:15–21.

Essner JJ. 2016. Zebrafish embryo microinjection: Ribonucleoprotein delivery using the Alt-RTM CRISPR-Cas9 System. In.

Essner JJ, Amack JD, Nyholm MK, Harris EB, Yost HJ. 2005. Kupffer’s vesicle is a ciliated organ of asymmetry in the zebrafish embryo that initiates left-right development of the brain, heart and gut. Development 132:1247 LP–1260.

Figiel DM, Elsayed R, Nelson AC. 2021. Investigating the molecular guts of endoderm formation using zebrafish. Brief Funct Genomics.

Frankenberg S, Renfree MB. 2013. On the origin of POU5F1. BMC Biol 11:56.

Gamse JT, Kuan Y-S, Macurak M, Brösamle C, Thisse B, Thisse C, Halpern ME. 2005. Directional asymmetry of the zebrafish epithalamus guides dorsoventral innervation of the midbrain target. Development 132:4869–4881.

Gamse JT, Thisse C, Thisse B, Halpern ME. 2003. The parapineal mediates left-right asymmetry in the zebrafish diencephalon. Development 130:1059–1068.

Gokey JJ, Ji Y, Tay HG, Litts B, Amack JD. 2016. Kupffer’s vesicle size threshold for robust left-right patterning of the zebrafish embryo. Dev Dyn 245:22–33.

Gubbay J, Collignon J, Koopman P, Capel B, Economou A, Münsterberg A, Vivian N, Goodfellow P, Lovell-Badge R. 1990. A gene mapping to the sex-determining region of the mouse Y chromosome is a member of a novel family of embryonically expressed genes. Nature 346:245–250.

Hamada H, Meno C, Watanabe D, Saijoh Y. 2002. Establishment of vertebrate left– right asymmetry. Nature Reviews Genetics 3:103–113.

Horne-Badovinac S, Rebagliati M, Stainier DYR. 2003. A Cellular Framework for Gut-Looping Morphogenesis in Zebrafish. Science 302:662–665.

Iwafuchi M, Cuesta I, Donahue G, Takenaka N, Osipovich AB, Magnuson MA, Roder H, Seeholzer SH, Santisteban P, Zaret KS. 2020. Gene network transitions in embryos depend upon interactions between a pioneer transcription factor and core histones. Nature Genetics 52:418–427.

Jumper J, Evans R, Pritzel A, Green T, Figurnov M, Ronneberger O, Tunyasuvunakool K, Bates R, Žídek A, Potapenko A, et al. 2021. Highly accurate protein structure prediction with AlphaFold. Nature 596:583–589.

Kanai-Azuma M, Kanai Y, Gad JM, Tajima Y, Taya C, Kurohmaru M, Sanai Y, Yonekawa H, Yazaki K, Tam PPL, et al. 2002. Depletion of definitive gut endoderm in Sox17-null mutant mice. Development.

Karlsson J, von Hofsten J, Olsson P-E. 2001. Generating Transparent Zebrafish: A Refined Method to Improve Detection of Gene Expression During Embryonic Development. Marine Biotechnology 3:522–527.

Kikuchi Y, Agathon A, Alexander J, Thisse C, Waldron S, Yelon D, Thisse B, Stainier DYR. 2001. casanova encodes a novel Sox-related protein necessary and sufficient for early endoderm formation in zebrafish. Genes and Development.

Kikuchi Y, Trinh LA, Reiter JF, Alexander J, Yelon D, Stainier DYR. 2000. The zebrafish bonnie and clyde gene encodes a Mix family homeodomain protein that regulates the generation of endodermal precursors. Genes and Development.

Kleinhans DS, Lecaudey V. 2019. Standardized mounting method of (zebrafish) embryos using a 3D-printed stamp for high-content, semi-automated confocal imaging. BMC Biotechnology 19:68.

Kondoh H, Kamachi Y. 2010. SOX-partner code for cell specification: Regulatory target selection and underlying molecular mechanisms. Int J Biochem Cell Biol 42:391–399.

Kramer-Zucker AG, Olale F, Haycraft CJ, Yoder BK, Schier AF, Drummond IA. 2005. Cilia-driven fluid flow in the zebrafish pronephros, brain and Kupffer&#039;s vesicle is required for normal organogenesis. Development 132:1907 LP–1921.

Kuleshov MV, Jones MR, Rouillard AD, Fernandez NF, Duan Q, Wang Z, Koplev S, Jenkins SL, Jagodnik KM, Lachmann A, et al. 2016. Enrichr: a comprehensive gene set enrichment analysis web server 2016 update. Nucleic Acids Research 44:W90–W97.

Labun K, Montague TG, Krause M, Torres Cleuren YN, Tjeldnes H, Valen E. 2019. CHOPCHOP v3: expanding the CRISPR web toolbox beyond genome editing. Nucleic Acids Research 47:W171–W174.

Lilly AJ, Lacaud G, Kouskoff V. 2017. SOXF transcription factors in cardiovascular development. Semin Cell Dev Biol 63:50–57.

Livak KJ, Schmittgen TD. 2001. Analysis of relative gene expression data using real-time quantitative PCR and the 2(-Delta Delta C(T)) Method. Methods (San Diego, Calif.) 25:402–408.

Livingstone CD, Barton GJ. 1993. Protein sequence alignments: a strategy for the hierarchical analysis of residue conservation. Computer applications in the biosciences : CABIOS 9:745–756.

Long S, Ahmad N, Rebagliati M. 2003. The zebrafish nodal-related gene southpaw is required for visceral and diencephalic left-right asymmetry. Development (Cambridge, England) 130:2303–2316.

Love MI, Huber W, Anders S. 2014. Moderated estimation of fold change and dispersion for RNA-seq data with DESeq2. Genome Biology 15:550.

Lunde K, Belting HG, Driever W. 2004. Zebrafish pou5f1/pou2, Homolog of Mammalian Oct4, Functions in the Endoderm Specification Cascade. Current Biology.

McCurley AT, Callard GV. 2008. Characterization of housekeeping genes in zebrafish: male-female differences and effects of tissue type, developmental stage and chemical treatment. BMC Molecular Biology 9:102.

McDonald Angela CH, Biechele S, Rossant J, Stanford William L. 2014. Sox17-Mediated XEN Cell Conversion Identifies Dynamic Networks Controlling Cell-Fate Decisions in Embryo-Derived Stem Cells. Cell Reports 9:780–793.

Melby AE, Warga RM, Kimmel CB. 1996. Specification of cell fates at the dorsal margin of the zebrafish gastrula. Development 122:2225 LP-2237.

Milewski WM, Duguay SJ, Chan SJ, Steiner DF. 1998. Conservation of PDX-1 Structure, Function, and Expression in Zebrafish*. Endocrinology 139:1440–1449.

Miller AZ, Satchie A, Tannenbaum AP, Nihal A, Thomson JA, Vereide DT. 2018. Expandable Arterial Endothelial Precursors from Human CD34+ Cells Differ in Their Proclivity to Undergo an Endothelial-to-Mesenchymal Transition. Stem Cell Reports 10:73–86.

Mizoguchi T, Verkade H, Heath JK, Kuroiwa A, Kikuchi Y. 2008. Sdf1/Cxcr4 signaling controls the dorsal migration of endodermal cells during zebrafish gastrulation. Development 135:2521–2529.

Montague TG, Gagnon JA, Schier AF. 2018. Conserved regulation of Nodal-mediated left-right patterning in zebrafish and mouse.

Mootha VK, Lindgren CM, Eriksson K-F, Subramanian A, Sihag S, Lehar J, Puigserver P, Carlsson E, Ridderstråle M, Laurila E, et al. 2003. PGC-1α-responsive genes involved in oxidative phosphorylation are coordinately downregulated in human diabetes. Nature Genetics 34:267–273.

Mukherjee S, Chaturvedi P, Rankin SA, Fish MB, Wlizla M, Paraiso KD, MacDonald M, Chen X, Weirauch MT, Blitz IL, et al. 2020. Sox17 and β-catenin co-occupy Wnt-responsive enhancers to govern the endoderm gene regulatory network. eLife 9:e58029–e58029.

Mukherjee S, Luedeke DM, McCoy L, Iwafuchi M, Zorn AM. 2022. SOX transcription factors direct TCF-independent WNT/&#x3b2;-catenin responsive transcription to govern cell fate in human pluripotent stem cells. Cell Reports 40.

Nelson AC, Cutty SJ, Niini M, Stemple DL, Flicek P, Houart C, Bruce AEE, Wardle FC. 2014. Global identification of smad2 and eomesodermin targets in zebrafish identifies a conserved transcriptional network in mesendoderm and a novel role for eomesodermin in repression of ectodermal gene expression. BMC Biology.

Ng CKL, Li NX, Chee S, Prabhakar S, Kolatkar PR, Jauch R. 2012. Deciphering the Sox-Oct partner code by quantitative cooperativity measurements. Nucleic Acids Research 40:4933–4941.

Niakan JHMR, Vokes SA, Rodolfa KT, Sherwood RI, Yamaki M, Dimos JT, Chen AE, Melton DA, McMahon AP, Eggan K. 2010. Sox17 promotes differentiation in mouse embryonic stem cells by directly regulating extraembryonic gene expression and indirectly antagonizing self-renewal. Genes and Development 24:312–326.

Odenthal J, Nüsslein-Volhard C. 1998. fork head domain genes in zebrafish. Development Genes and Evolution 208:245–258.

Palasingam P, Jauch R, Ng CKL, Kolatkar PR. 2009. The structure of Sox17 bound to DNA reveals a conserved bending topology but selective protein interaction platforms. Journal of molecular biology 388:619–630.

Pani L, Overdier DG, Porcella A, Qian X, Lai E, Costa RH. 1992. Hepatocyte Nuclear Factor 3β Contains Two Transcriptional Activation Domains, One of Which Is Novel and Conserved with the Drosophila Fork Head Protein. Molecular and Cellular Biology 12:3723–3732.

Pevny LH, Lovell-Badge R. 1997. Sox genes find their feet. Current opinion in genetics & development 7:338–344.

Poulain M. 2006. Zebrafish endoderm formation is regulated by combinatorial Nodal, FGF and BMP signalling. Development.

Rebagliati MR, Toyama R, Fricke C, Haffter P, Dawid IB. 1998. Zebrafish Nodal-Related Genes Are Implicated in Axial Patterning and Establishing Left–Right Asymmetry. Developmental Biology 199:261–272.

Reim G, Mizoguchi T, Stainier DY, Kikuchi Y, Brand M. 2004. The POU domain protein Spg (Pou2/Oct4) is essential for endoderm formation in cooperation with the HMG domain protein casanova. Developmental Cell.

Saba R, Kitajima K, Rainbow L, Engert S, Uemura M, Ishida H, Kokkinopoulos I, Shintani Y, Miyagawa S, Kanai Y, et al. 2019. Endocardium differentiation through Sox17 expression in endocardium precursor cells regulates heart development in mice. Scientific Reports 9:11953.

Sarkar A, Hochedlinger K. 2013. The sox family of transcription factors: versatile regulators of stem and progenitor cell fate. Cell stem cell 12:15–30.

Saund RS, Kanai-Azuma M, Kanai Y, Kim I, Lucero MT, Saijoh Y. 2012. Gut endoderm is involved in the transfer of left-right asymmetry from the node to the lateral plate mesoderm in the mouse embryo. Development 139:2426–2435.

Schneider I, Houston DW, Rebagliati MR, Slusarski DC. 2008. Calcium fluxes in dorsal forerunner cells antagonize β-catenin and alter left-right patterning. Development 135:75–84.

Séguin CA, Draper JS, Nagy A, Rossant J. 2008. Establishment of Endoderm Progenitors by SOX Transcription Factor Expression in Human Embryonic Stem Cells. Cell stem cell 3.

Sievers F, Higgins DG. 2018. Clustal Omega for making accurate alignments of many protein sequences. Protein Science 27:135–145.

Sievers F, Wilm A, Dineen D, Gibson TJ, Karplus K, Li W, Lopez R, McWilliam H, Remmert M, Söding J, et al. 2011. Fast, scalable generation of high-quality protein multiple sequence alignments using Clustal Omega. Molecular Systems Biology 7:539.

Sinclair AH, Berta P, Palmer MS, Hawkins JR, Griffiths BL, Smith MJ, Foster JW, Frischauf AM, Lovell-Badge R, Goodfellow PN. 1990. A gene from the human sex-determining region encodes a protein with homology to a conserved DNA-binding motif. Nature 346:240–244.

Sinner D, Rankin S, Lee M, Zorn AM. 2004. Sox17 and beta-catenin cooperate to regulate the transcription of endodermal genes. Development 131:3069–3080.

Stafford D, White RJ, Kinkel MD, Linville A, Schilling TF, Prince VE. 2006. Retinoids signal directly to zebrafish endoderm to specify insulin-expressing β-cells. Development 133:949–956.

Stefanovic S, Abboud N, Desilets S, Nury D, Cowan C, Puceat M. 2009. Interplay of Oct4 with Sox2 and Sox17: a molecular switch from stem cell pluripotency to specifying a cardiac fate. J Cell Biol 186:665–673.

Subramanian A, Tamayo P, Mootha VK, Mukherjee S, Ebert BL, Gillette MA, Paulovich A, Pomeroy SL, Golub TR, Lander ES, et al. 2005. Gene set enrichment analysis: A knowledge-based approach for interpreting genome-wide expression profiles. Proceedings of the National Academy of Sciences 102:15545–15550.

Sutherland MJ, Ware SM. 2009. Disorders of left–right asymmetry: Heterotaxy and situs inversus. American Journal of Medical Genetics Part C: Seminars in Medical Genetics 151C:307-317.

Sutherland RJ. 1982. The dorsal diencephalic conduction system: A review of the anatomy and functions of the habenular complex. Neuroscience & Biobehavioral Reviews 6:1–13.

Talbot CD, Walsh MD, Cutty SJ, Elsayed R, Vlachaki E, Bruce AEE, Wardle FC, Nelson AC. 2022. Eomes function is conserved between zebrafish and mouse and controls left-right organiser progenitor gene expression via interlocking feedforward loops. Frontiers in Cell and Developmental Biology 10.

Tasnim M, Wahlquist P, Hill JT. 2024. Zebrafish: unraveling genetic complexity through duplicated genes. Dev Genes Evol.

The UniProt C. 2023. UniProt: the Universal Protein Knowledgebase in 2023. Nucleic Acids Research 51:D523–D531.

Thisse B, Pflumio, S., Fürthauer, M., Loppin, B., Heyer, V., Degrave, A., Woehl, R., Lux, A., Steffan, T., Charbonnier, X.Q. and Thisse, C. (2001). 2001. Expression of the zebrafish genome during embryogenesis (NIH R01 RR15402). In. ZFIN Direct Data Submission.

Thisse C, Thisse B. 2008. High-resolution in situ hybridization to whole-mount zebrafish embryos. Nature Protocols 3:59–69.

Traver D, Paw BH, Poss KD, Penberthy WT, Lin S, Zon LI. 2003. Transplantation and in vivo imaging of multilineage engraftment in zebrafish bloodless mutants. Nature Immunology 4:1238–1246.

Varadi M, Anyango S, Deshpande M, Nair S, Natassia C, Yordanova G, Yuan D, Stroe O, Wood G, Laydon A, et al. 2022. AlphaFold Protein Structure Database: massively expanding the structural coverage of protein-sequence space with high-accuracy models. Nucleic Acids Research 50:D439–D444.

Viotti M, Niu L, Shi S-H, Hadjantonakis A-K. 2012. Role of the Gut Endoderm in Relaying Left-Right Patterning in Mice. PLOS Biology 10:e1001276–e1001276.

Voldoire E, Brunet F, Naville M, Volff JN, Galiana D. 2017. Expansion by whole genome duplication and evolution of the sox gene family in teleost fish. PLoS ONE 12:1–20.

Wagner DE, Weinreb C, Collins ZM, Briggs JA, Megason SG, Klein AM. 2018. Single-cell mapping of gene expression landscapes and lineage in the zebrafish embryo. Science 360:981 LP-987.

Warga RM, Kane DA. 2018. Wilson cell origin for kupffer’s vesicle in the zebrafish. Developmental Dynamics 247:1057–1069.

Wegner M. 2010. All purpose Sox: The many roles of Sox proteins in gene expression. The International Journal of Biochemistry & Cell Biology 42:381–390.

Westerfield M. 2000. The Zebrafish Book. A Guide for The Laboratory Use of Zebrafish (Danio rerio). Eugene 385.

Wu RS, Lam II, Clay H, Duong DN, Deo RC, Coughlin SR. 2018. A Rapid Method for Directed Gene Knockout for Screening in G0 Zebrafish. Developmental Cell 46:112–125.e114.

Yelon D, Horne SA, Stainier DYR. 1999. Restricted Expression of Cardiac Myosin Genes Reveals Regulated Aspects of Heart Tube Assembly in Zebrafish. Developmental Biology 214:23–37.

Yuan H, Corbi N, Basilico C, Dailey L. 1995. Developmental-specific activity of the FGF-4 enhancer requires the synergistic action of Sox2 and Oct-3. Genes and Development 9:2635–2645.

Zhao J, Lambert G, Meijer AH, Rosa FM. 2013. The transcription factor Vox represses endoderm development by interacting with x and Pou2. Development.

Zhong TP, Childs S, Leu JP, Fishman MC. 2001. Gridlock signalling pathway fashions the first embryonic artery. Nature 414:216–220.

