## Supplementary Material and Tables for "The molecular basis for functional divergence of duplicated SOX factors controlling endoderm formation and left-right patterning in zebrafish"

Supplementary Figures and Tables

Supplementary Figures

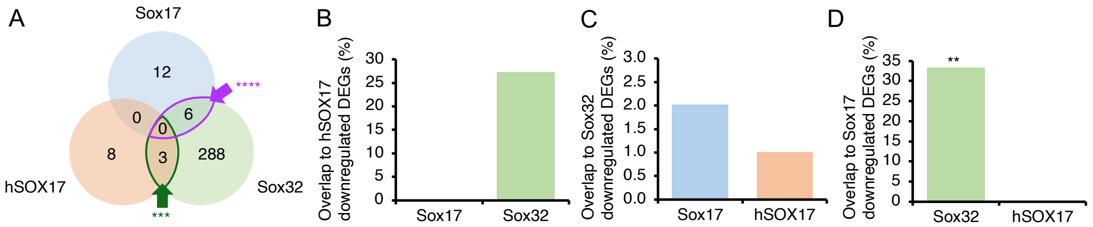


**Supplementary Figure 1: Downregulated differentially expressed genes (DEGs) analysis for hSOX17, Sox32 and Sox17 RNA-seq**. A) Venn diagram indicating overlap in genes downregulated by overexpression of each of the TFs. Bar charts indicating percentage overlap of significantly downregulated genes on hSOX17 OE (B), Sox32 OE (C), and Sox17 OE (D) to those downregulated by the other TFs. Bars represent the percentage of genes downregulated by the factor on the y-axis also downregulated by the factor on the x-axis. Statistical tests to determine whether the overlap was significantly greater with one factor on the x-axis compared to the other were carried out using Fisher's Exact test. Note that the only significant overlap (as determined using Fisher's Exact test) is between genes downregulated by Sox32 and Sox17. ** P<0.01.

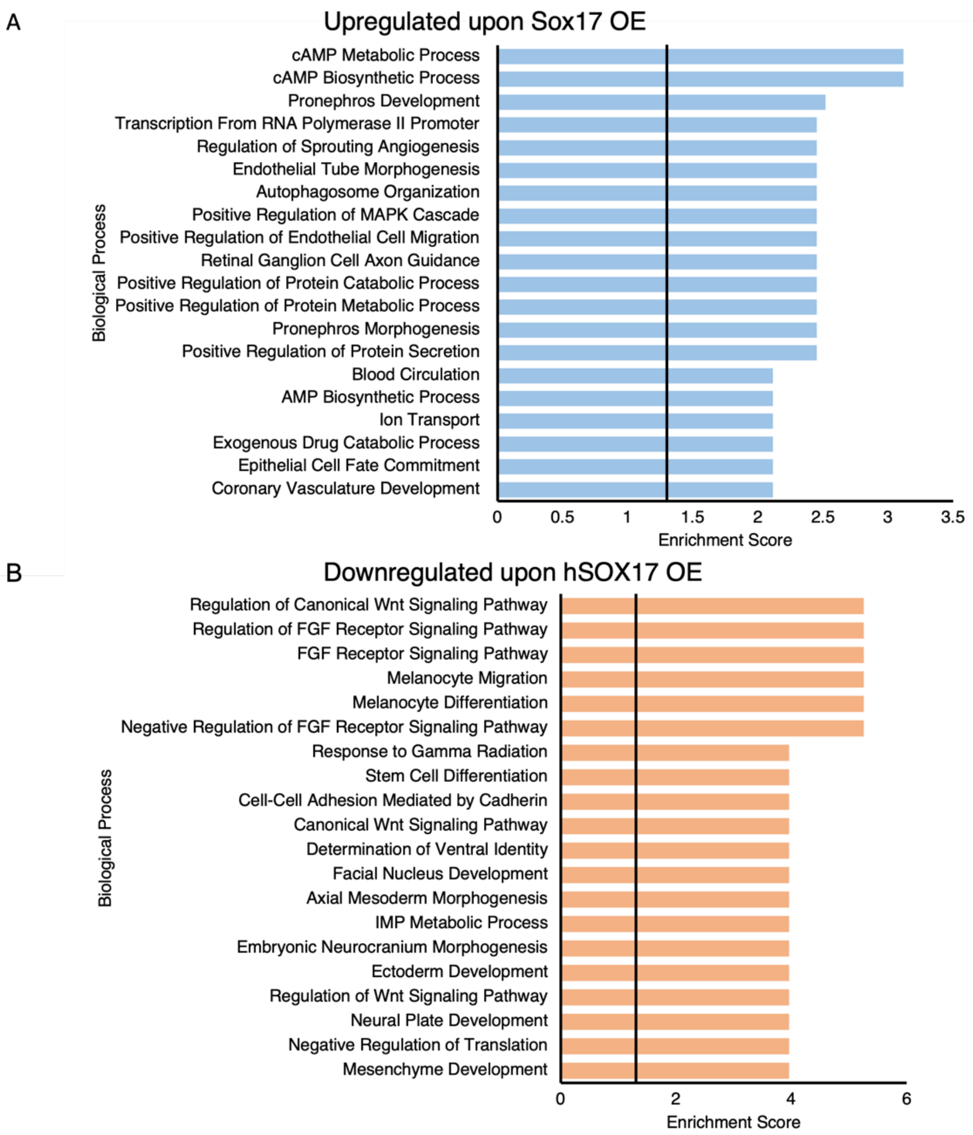


**Supplementary Figure 2: Functional annotation analysis carried out using fishEnrichR for Sox17 (A) and hSOX17(B).** A) GO analysis carried out on upregulated DEGs due to Sox17 OE. B) GO analysis carried out on downregulated DEGs due to hSOX17 OE. FDR<0.05 cut off applied with top 20 biological process terms graphed. Vertical lines indicate enrichment score from FDR = 0.05

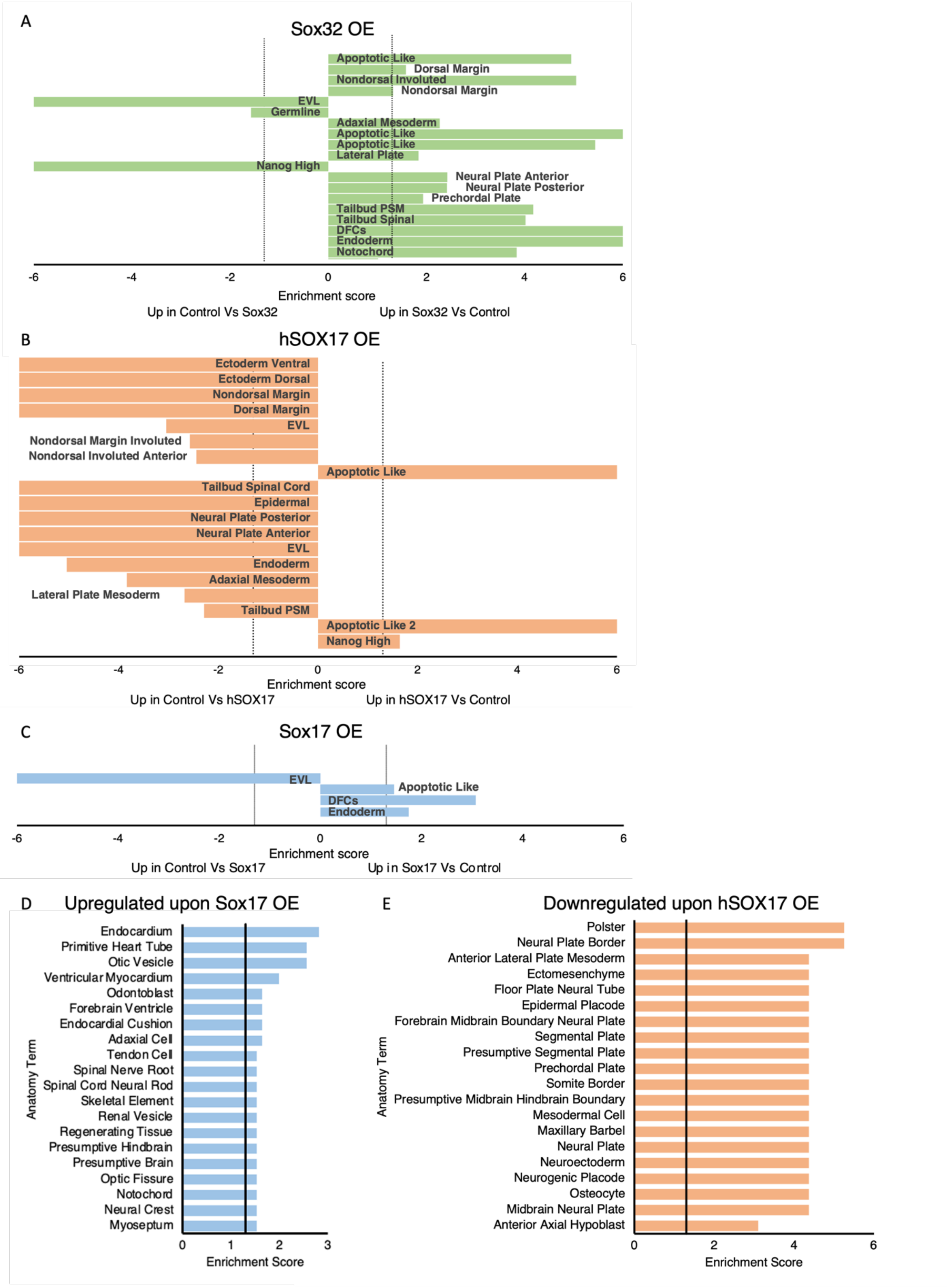
A
Sox32 OE
Apoptotic Like
Dorsal Margin
Nondorsal Involuted
Nondorsal Margin
EVL
Germline
Adaxial Mesoderm
Apoptotic Like
Apoptotic Like
Lateral Plate
Nanog High
Neural Plate Anterior
Neural Plate Posterior
Prechordal Plate
Tailbud PSM
Tailbud Spinal
DFCs
Endoderm
Notochord
-6 -4 -2 0 2 4 6
Enrichment score
Up in Control Vs Sox32
Up in Sox32 Vs Control
B
hSOX17 OE
Ectoderm Ventral
Ectoderm Dorsal
Nondorsal Margin
Dorsal Margin
EVL
Nondorsal Margin Involuted
Nondorsal Involuted Anterior
Apoptotic Like
Tailbud Spinal Cord
Epidermal
Neural Plate Posterior
Neural Plate Anterior
EVL
Endoderm
Adaxial Mesoderm
Lateral Plate Mesoderm
Tailbud PSM
Apoptotic Like 2
Nanog High
-6 -4 -2 0 2 4 6
Enrichment score
Up in Control Vs hSOX17
Up in hSOX17 Vs Control
C
Sox17 OE
EVL
Apoptotic Like
DFCs
Endoderm
-6 -4 -2 0 2 4 6
Enrichment score
Up in Control Vs Sox17
Up in Sox17 Vs Control
D
Upregulated upon Sox17 OE
Endocardium
Primitive Heart Tube
Otic Vesicle
Ventricular Myocardium
Odontoblast
Forebrain Ventricle
Endocardial Cushion
Adaxial Cell
Tendon Cell
Spinal Nerve Root
Spinal Cord Neural Rod
Skeletal Element
Renal Vesicle
Regenerating Tissue
Presumptive Hindbrain
Presumptive Brain
Optic Fissure
Notochord
Neural Crest
Myoseptum
Anatomy Term
0 1 2 3
Enrichment Score
E
Downregulated upon hSOX17 OE
Polster
Neural Plate Border
Anterior Lateral Plate Mesoderm
Ectomesenchyme
Floor Plate Neural Tube
Epidermal Placode
Forebrain Midbrain Boundary Neural Plate
Segmental Plate
Presumptive Segmental Plate
Prechordal Plate
Somite Border
Presumptive Midbrain Hindbrain Boundary
Mesodermal Cell
Maxillary Barbel
Neural Plate
Neuroectoderm
Neurogenic Placode
Osteocyte
Midbrain Neural Plate
Anterior Axial Hypoblast
Anatomy Term
0 2 4 6
Enrichment Score

**Supplementary Figure 3: Anatomical enrichment analysis**. A-C) Gene set enrichment analysis (GSEA) carried out against cell type markers defined by scRNA-seq at 6-8 hpf (Wagner et al., 2018). Enrichment score was calculated through -$\log_{10}$(FDR q-value) with a cut off of FDR q-value <0.05 applied. Vertical lines indicate enrichment score from FDR = 0.05. EVL = enveloping layer, DFC= dorsal forerunner cell, PSM = presomitic mesoderm, nondorsal involuted anterior includes the presumptive endoderm. A) Sox32 upregulated transcripts show enrichment for 8 hpf endoderm markers as expected. While 8 hpf adaxial mesoderm markers are enriched amongst Sox32-induced transcripts, 46% of genes that contribute to this enrichment are also 8 hpf endoderm markers within the scRNA-seq dataset. Removal of these overlapping genes from analysis results in no enrichment of the adaxial mesoderm by Sox32 (data not shown). hSOX17 does not show enrichment for endoderm markers (B). Sox17 overexpression shows induction of 8 hpf endoderm markers but at a much weaker levels compared to Sox32 and lacks enrichment for markers of “nondorsal involuted anterior” as defined by Wagner et al., which includes the presumptive endoderm. D-E) Anatomy term enrichment analysis carried out using fishEnrichR for genes significantly upregulated by Sox17 (D) and genes significantly downregulated by hSOX17(E).

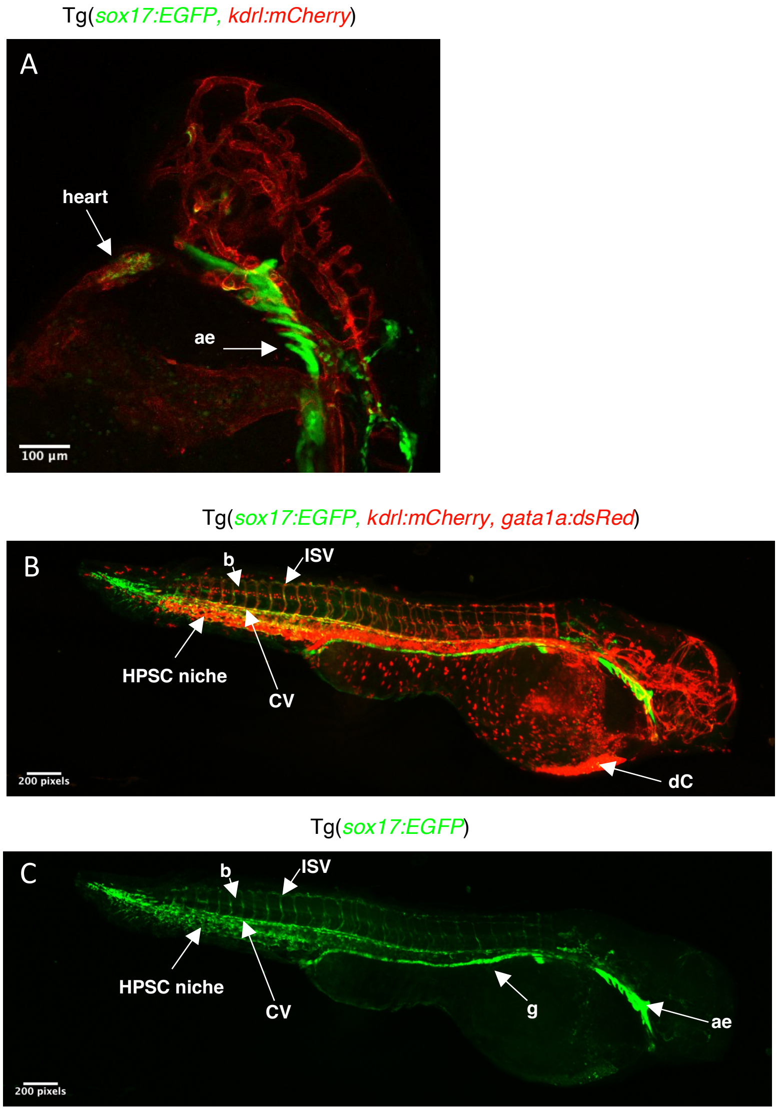


**Supplementary Figure 4: Expression of sox17:EGFP in the heart and vasculature.** Lateral images of 48hpf heterozygous *sox17:EGFP* and *kdrl:mCherry* embryos **(A)** and triple transgenic embryos heterozygous to *sox17:EGFP*, *kdrl:mCherry* and *gata1a:dsRed* **(B-C)**. **(A)** *sox17:EGFP* exhibits co-expression with *kdrl:mCherry* in the heart, with *kdrl:mCherry* marking the endocardium specifically. A) Maximum Z projection of all slices. **(B)** *sox17:EGFP* exhibits co-localisation with endothelial marker *kdrl:mCherry* in the intersomitic vessels (ISV) and cardinal vein (CV), with a lack of co-expression in the duct of Cuvier (dC). Additional *sox17:EGFP* co-expression is observed with erythroid marker *gata1a:dsRed* in the blood (b) and haemopoietic stem and progenitor cell (HPSC) niche.**(C)** Identical image to C but presenting *sox17:EGFP* expression alone to show presence within endodermal, endothelial and erythroid lineages. ae = anterior endoderm, HSPC = haemopoietic stem and progenitor cell niche, b = blood, CV = cardinal vein, ISV = intersomitic vessels, dC = duct of Curvier, g = gut

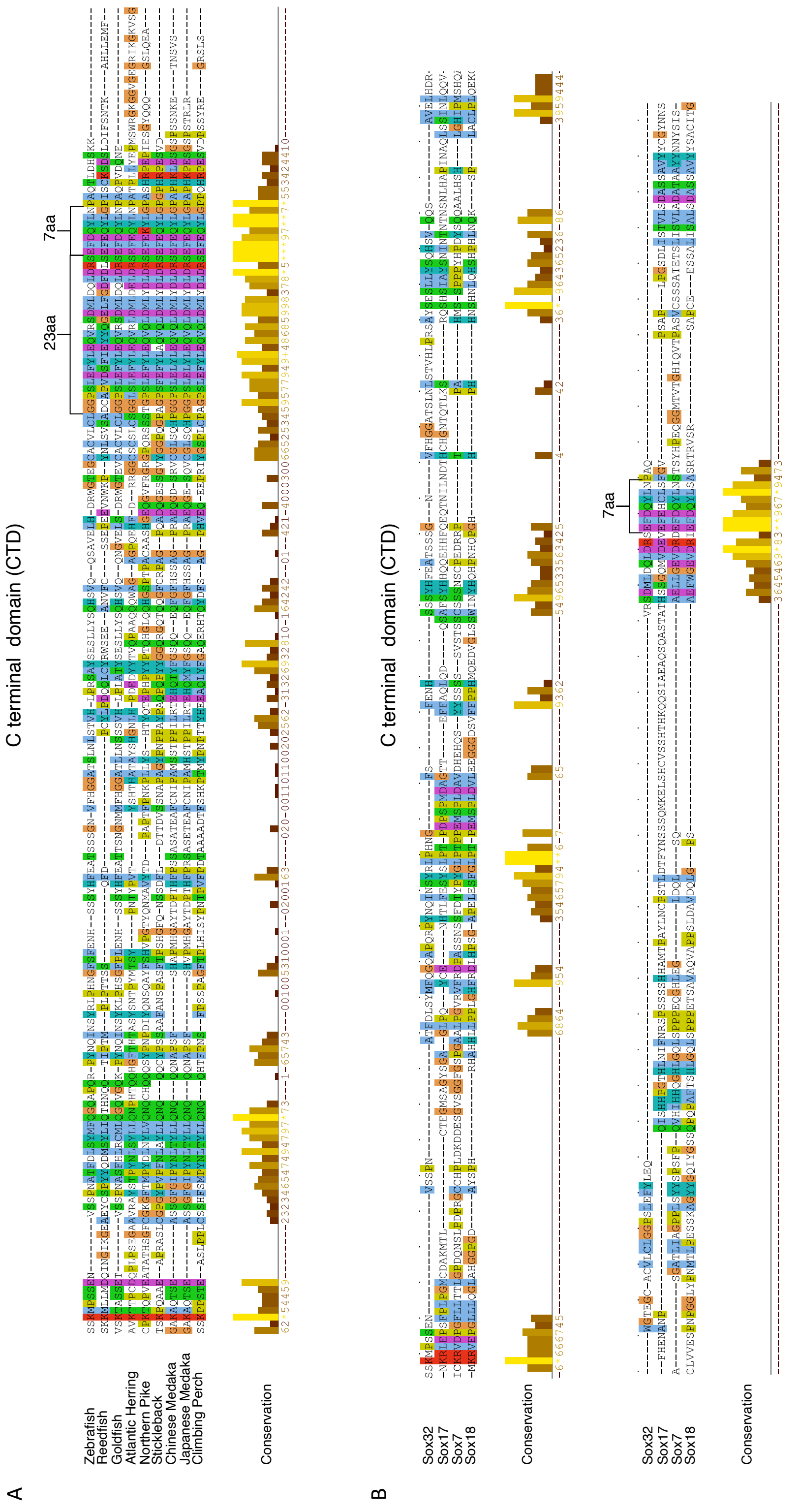


**Supplementary Figure 5: Alignments of Sox32 C terminal domain with Sox32 orthologues (A) and zebrafish SoxF subfamily proteins (B).** Alignments conducted using ClustalOmega in JalView and colour coded based on conserved amino acid properties. Conservation levels calculated according to Analysis of Multiply Aligned Sequences (AMAS) are displayed. A) Annotation of highly conserved 23 aa, plus the 7 aa peptide within the CTD previously identified by Kikuchi et al. (2001) and Sinner et al., (2004), B) Annotation of 7 aa peptide (equivalent to 7 aa domain in A) within the CTD. Indicates a high degree of conservation between Sox32, Sox7 and Sox18, while Sox17 shows divergence.

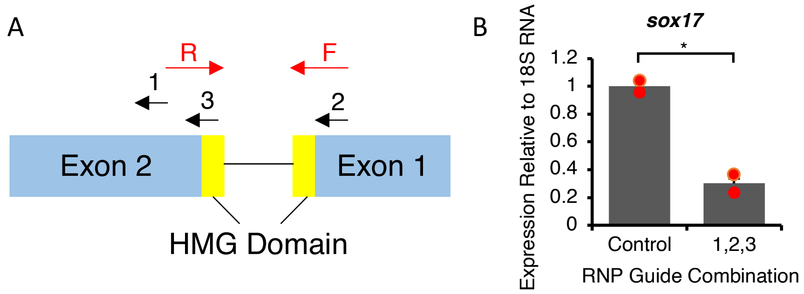


**Supplementary Figure 6: Guide RNAs against *sox17* efficiently disrupt the gene locus A)** Gene model representing the exons and HMG domain of Sox17 with gRNA positioning (1-3) and qPCR primers (F,R) annotated**. B)** *sox17* qPCR normalised to 18S rRNA on 75% epiboly (8 hpf) control or *sox17* Cas9 RNP injected embryos with indicated gRNAs. Statistically significant differences were inferred using Students t-test test on independent biological duplicate datasets, * $P < 0.05$. Error bars represent relative standard deviation.

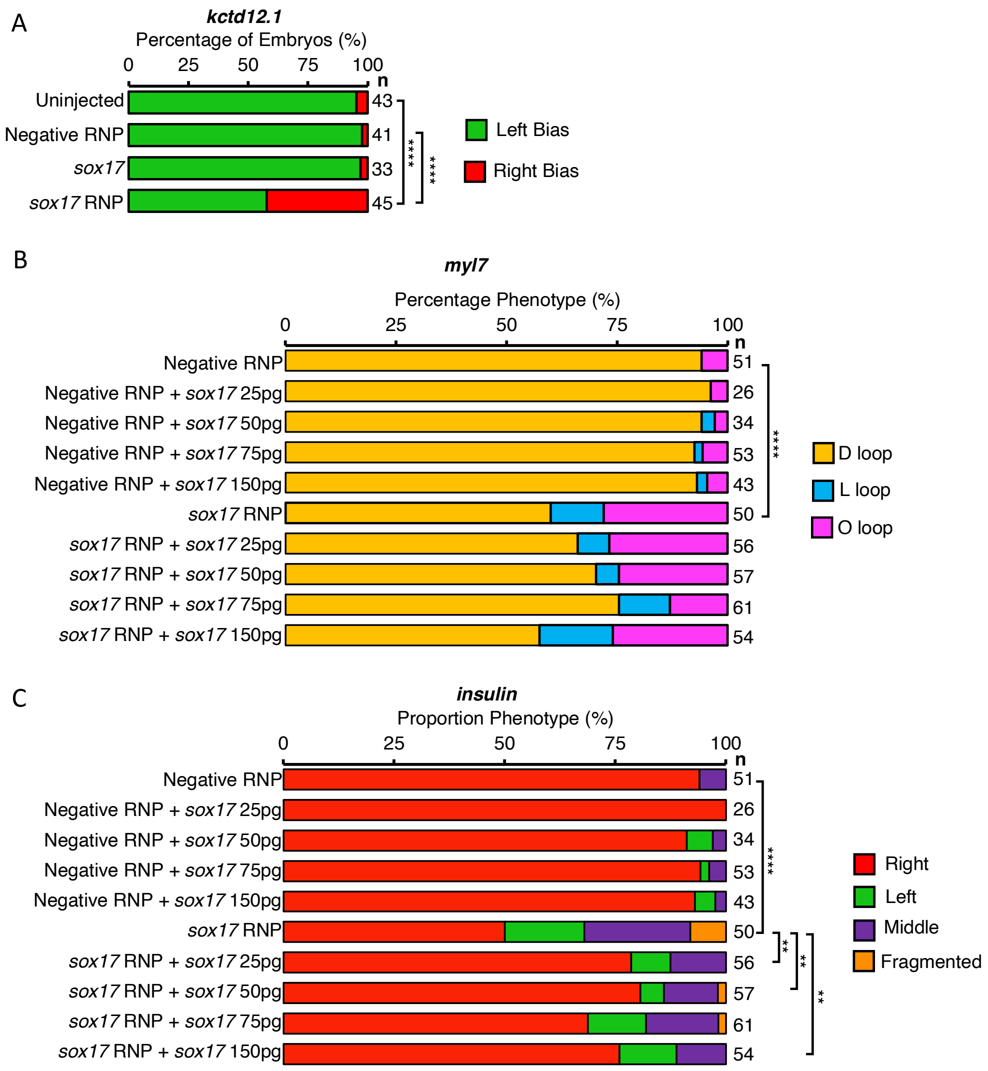


**Supplementary Figure 7: Sox17 overexpression does not affect organ asymmetry. A**) Whole mount *in situ* hybridisation (WISH) for asymmetrically expressed habenula marker *kctd12.1* carried out at 4 dpf. Statistically significant differences in categorical scoring was inferred using Fisher's Exact test on independent biological triplicate datasets. **B)** WISH for cardiac marker *myl7* at 48 hpf. Experiment carried out on a range of conditions to investigate dose dependent overexpression of Sox17 and rescue of Sox17 RNP phenotype by Sox17 mRNA. **C)** WISH for endocrine pancreas marker *insulin* at 48 hpf. Experiment carried out on a range of conditions to investigate dose dependent overexpression of Sox17 and rescue of Sox17 RNP phenotype by Sox17 mRNA. Statistically significant differences in categorical scoring was inferred using Fisher's Exact test on independent biological duplicate datasets.** P<0.01, **** P<0.0001. n = number of embryos analysed. Representative images can be viewed in Figure 6.

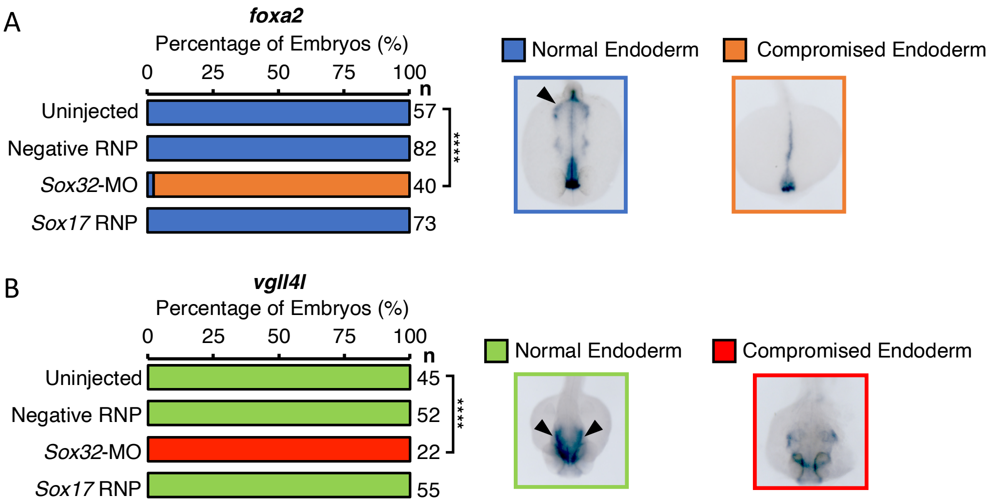


**Supplementary Figure 8: Anterior endoderm is present in Sox17 CRISPants.** Whole mount *in situ* hybridisation for anterior endoderm in uninjected, Negative RNP, *sox32* MO (positive control) and *sox17* RNP injected embryos. **A)** *foxa2* at 24 hpf. **B)** *vgll4l* at 48 hpf. Dorsal views with anterior to the bottom, arrowheads indicate anterior endoderm. Statistically significant differences in categorical scoring was inferred using Fisher's Exact test on independent biological triplicate datasets, **** P < 0.0001. Sox32 MO injected embryos showed compromised, absent anterior endoderm as expected, while anterior endoderm appears normal in Sox17 CRISPants. n = total number of embryos analysed.

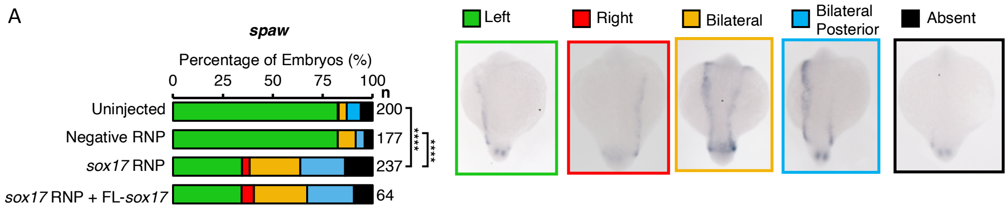


**Supplementary Figure 9: Sox17 CRISPants exhibit abnormal *spaw* expression that cannot be rescued through co-injection of exogenous *sox17* mRNA.** Wholemount *in situ* hybridisation for *spaw* conducted at 18 somite stage. Sox17 CRISPants show a significant increase in the percentage of embryos depicting abnormal *spaw* expression, phenotypes include right LPM expression, bilateral, bilateral posterior and absent expression. Abnormal *spaw* expression could not be rescued by co-injection of sox17 RNP with exogenous sox17 mRNA. Statistically significant differences in categorical scoring was inferred using Fisher's Exact test on independent biological triplicate datasets, **** P<0.0001. n= number of embryos analysed.

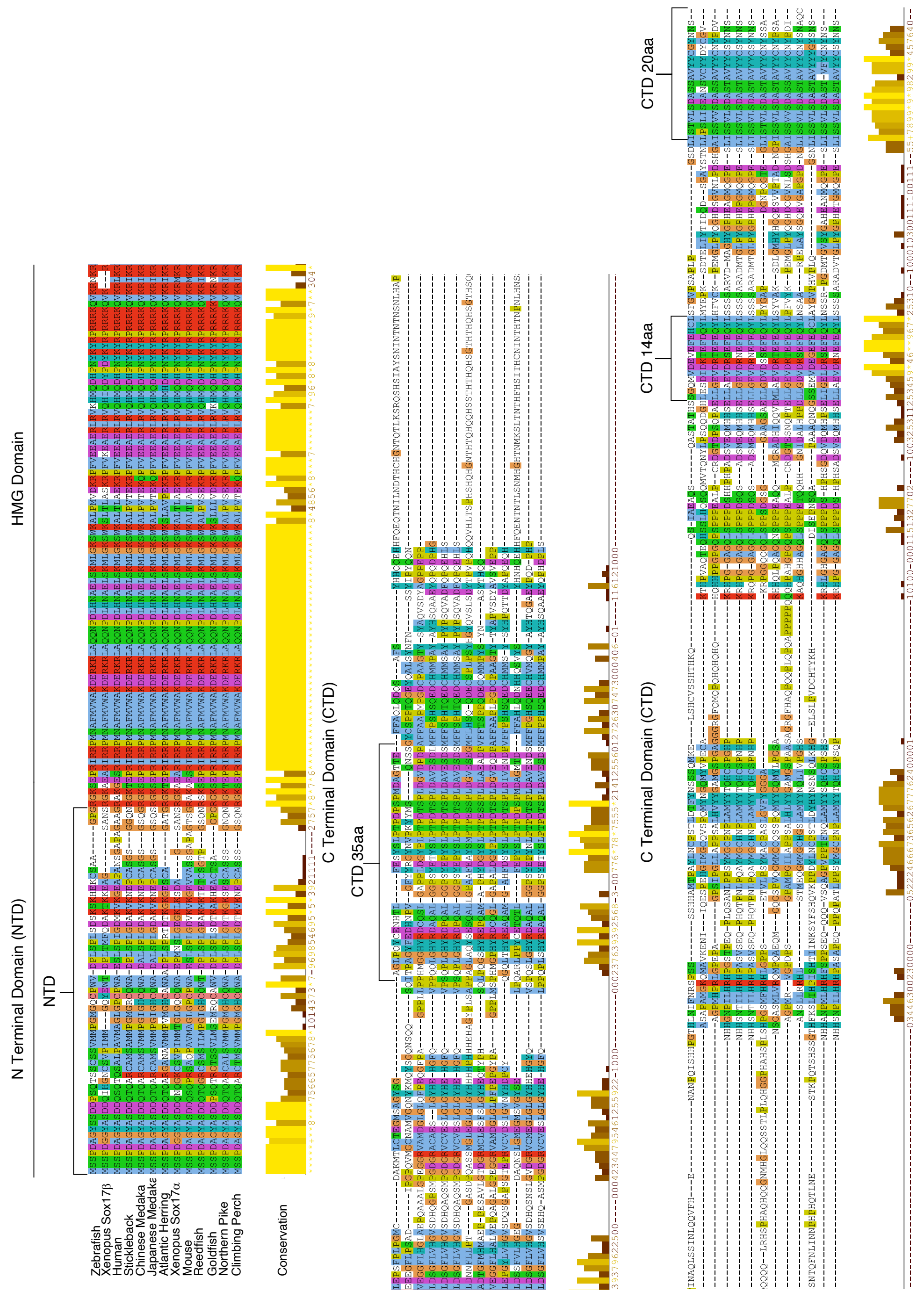


**Supplementary Figure 10: Alignment of Sox17 teleost and mammalian orthologues.** Alignment carried out using Clustal Omega in Jalview and colour coded based on conserved amino acid properties. Conservation levels calculated according to Analysis of Multiply Aligned Sequences (AMAS) are displayed. Four distinct highly conserved domains predicted to be important for the role of Sox17 in LR patterning selected for deletion are indicated in brackets.

**Supplementary Video 1 - Co-localisation of *sox17:EGFP* and *kdrl:mCherry* expression in the zebrafish heart.** Lateral view of the heart region shown in Figure S4A. 48hpf embryos heterozygous for *sox17:EGFP* and *kdrl:mCherry*. *kdrl:mCherry* marks the vascular endothelial lineage. Video depicting all Z slices from heart region shown in Figure S4A.

### Supplementary Tables

**Supplementary Table 11: Deletion and hybrid constructs.** Depicts the amino acids removed or added to Sox32/Sox17 to generate constructs referred to within the manuscript.

| pCS2+ Construct | Description |
|---|---|
| Sox17Δ7aa | Removal of amino acids 374-380 (EFEHCLS) |
| Sox32Δ7aa | Removal of amino acids 291-297 (EFDQYLN) |
| Sox17Δ7aa+Sox32 7aa | Replacement of Sox17 amino acids 374-380 (EFEHCLS) with Sox32 amino acids 291-297 (EFDQYLN) |
| Sox32Δ7aa+Sox17 7aa | Replacement of Sox32 amino acids 291-297 (EFDQYLN) with Sox17 amino acids 374-380 (EFEHCLS) |
| Sox17Δ7aa+Sox32 25aa | Replacement of Sox17 amino acids 374-380 (EFEHCLS) with Sox32 amino acids 373-397 (EFYLEQVRSDMLDQLDRSEFDQYLN) |
| Sox17ΔNTD | Removal of amino acids 1-55 (MSSPDAGYSSDDPSQTSSCSSVMMPGMGQCPW VDPLSPLSDSKSKHEKCSAAGPG) |
| Sox17ΔCTD35aa | Removal of amino acids 173-207 (GAGLPQYCENHTLFESYSLPTPDPSPMDAGTTEFF ) |
| Sox17ΔCTD14aa | Removal of amino acids 366-379 (SGQMVDEVEFEHCL) |
| Sox17ΔCTD20aa | Removal of amino acids 394-413 (ISTVLSDASSAVYYCGYNNS) |
| Sox32ΔHMG | Removal of amino acids 67-149(ETRVRRPLNAFIIWTKEERRRLAQLNPDLENTD LSKILGKTWKAMSLADKRPYMQEAERLRIQHTIDYP NYKYRPRRRKCNKR) |
| Sox32ΔHMG+ Sox17 HMG | Replacement of Sox32 amino acids 67-149 (ETRVRRPLNAFIIWTKEERRRLAQLNPDLENTDLSK ILGKTWKAMSLADKRPYMQEAERLRIQHTIDYPNYK YRPRRRKCNKR) with Sox17 amino acids 60-142 (EPRIRRPMNAFMVWAKDERKRLAQQNPDLHNAEL SKMLGKSWKALPMVDKRPFVEEAERLRVKHMQDH PNYKYRPRRRKQVKR) |

**Supplementary Table 12: Primers used for PCR to generate deletion and hybrid constructs.**

| pCS2+ Construct | F Primer (5’ → 3’) | R Primer (5’ → 3’) |
|---|---|---|
| myc-Sox32 | TCAGAAGAGGATCTGTATCTCGACCGGATGCTCCC | GATGAGTTTTTGTTCCATGCTGTTTTGCGTCCACT |
| myc-Sox32ΔNTD | CGAGTAAGACGCCCTTTAAA | CAGATCCTCTTCTGAGATGA |
| myc-Sox32ΔHMG | TGCAGCAAGATGCCTTCAAGTGAGAACGTCAGCTCTCCAAATGCCACCTTTGATC | CACGGGCGCTTTCGCTTCGGGACTTGAGCAGCTGGATTCAGACCCGACAGACAC |
| myc-Sox32ΔCTD23aa | GAATTTGACCAGTACCTCAATCC | CAAACACAGCACACACGCAC |
| myc-Sox32ΔCTD7aa | CCAGCACAGACTTTGGACCACAGC (Zhao et al., 2013) | ACTGCGATCAAGCTGGTCCAAC (Zhao et al., 2013) |
| Sox17Δ7 aa | TTTGGGGTCCCCAGTG | CACCTCGTCCACCATTTG |
| myc-Sox32Δ7aa + Sox17 7aa | CTGTCTCTCTCCAGCACAGACTTTGGAC | TGCTCAAACTCACTGCGATCAAGCTGGTC |
| Sox17Δ7aa + Sox32 7aa | AGTACCTCAATTTTGGGGTCCCCAGTG | GGTCAAATTCCACCTCGTCCACCATTTG |
| hSOX17 | CCGGCTCGAGGCCACCATGAGCAGCCCGGATG | CTAGTCTAGATTACACGTCAGGATAGTTGCA |
| HiFi primers for Sox17 HMG | AAGCGAAAGCGCCCGTGGAGCCGCGCATCCGAAGG | GAAGGCATCTTGCTGCATCGTTTCACCTGTTTGCGACGs |
| HiFi primers for Sox17Δ7aa+Sox32 25aa gblock | CCATCGATTCGAATTCAAGGATGAGCAGTCCCGATG | AGTTCTAGAGGCTCGAGAGGTCAAGAATTATTATAGCCGC |
| Sox17ΔNTD | CGCGGAAAGAGCGAG | CATCCTTGAATTCGAATCGA |
| Sox17ΔCTD35aa | GCCCAGCTTCAGGATC | TGAATAACCAGCACTCATAC |
| Sox17ΔCTD14aa | TTTGGGGTCCCCAGTGC | AGAACTGTGGGTTGCTGTAG |
| Sox17ΔCTD20aa | CCTCTCGAGCCTCTAGAACT | TAAGTCAGACCCCGGCAAAG |

**Supplementary Table 13: crRNA sequences used to generate guide RNAs for RNP complex production.**

| | crRNA (5' → 3') |
|---|---|
| Dr.Cas9.SOX17.1.AF | TACACCTGACCCCTCT CCTA |
| Dr.Cas9.SOX17.1.AE | ACACGAGAAATGCTCT GCGG |
| Dr.Cas9.SOX17.1.AC | GGACAAACGTCCATTC GTTG |
| Alt-R® CRISPR-Cas9 Negative Control crRNA #1 (cat# 1072544) | CGTTAATCGCGTATAAT ACG |
| Alt-R® CRISPR-Cas9 Negative Control crRNA #2 (cat# 1072545) | CATATTGCGCGTATAGT CGC |
| Alt-R® CRISPR-Cas9 Negative Control crRNA #3 (cat# 1072546)8 | GGCGCGTATAGTCGCG CGTA |

**Supplementary Table 14: qPCR primers.**

| | **F Primer (5'→3')** | **R Primer (5'→3')** |
|---|---|---|
| *18S* | TCGCTAGTTGGCATCGTTTA TG | CGGAGGTTCGAAGACGATCA |
| *fabp2* | GCCCATGACAACCTGAAGAT | TGTCCTTGCGTGTGAAAGTC |
| *tbx15* | GGTCAGCTTTGATAAACTCA AAC | CTGTTGGTTCTGGTAGGCT |
| *mcamb* | CTGCTATGCACAAGGCTACC | TTGGCAATGAGATCAGAGGTG |
| *foxj1a* | CAGATCCCACCTGGCAGAA CTC | ACTGGAGGTAATCTGCGCTTC |
| *pltp* | ACAACAGAGGAAACACTTG GACC | GGTCTCTGCCTTCAACATCTCC |
| *vgll4l* | CAGGACTGCTGCAATCACTC | GAAGGTTGGACTGCTTGGTG |
| *ndr1* | GCGAGCTGAACTTCGCATT | TCAATTAGCCCAACCGCAAG |
| *sox17* | CACAATGCGGAGCTGAGTAA | ATCGCTTGTTTCGTTTCACC |
| *cxcr4a* | TGGCTTATTACGAACACATC G | GAGCCGAATTCAGAGCTGTT |

**Supplementary Table 15: Species and accession numbers of teleost and mammalian Sox17 orthologues investigated for conservation.**

| Species | Accession Number |
|---|---|
| *Xenopus laevis* Sox17β.1 | O42601 |
| *Homo sapien* SOX17 | Q9H6I2 |
| *Danio rerio* Sox17 | Q5PQZ5 |
| *Gasterosteus aculeatus* Sox17 | G3NK71 |
| *Oryzias melastigma* Sox17 | A0A3B3B6G2 |
| *Oryzias sinensis* Sox17 | A0A8C7YKM0 |
| *Oryzias latipes* Sox17 | C3VV17 |
| *Takifugu rubripes* Sox17 | Q6WNS5 |
| *Tetraodon nigroviridis* Sox17 | H3DIC0 |
| *Cyprinus carpio* Sox17 | A0A9Q9WK15 |
| *Clupea harengus* Sox17 | A0A6P3VND2 |
| *Salmo salar* Sox17α | A0A1S3NCY8 |
| *Monopterus albus* Sox17 | Q8JGN3 |
| *Xenopus tropicalis* Sox17α | Q8AWH3 |
| *Mus musculus* SOX17 | Q61473-1 |
