## Supplementary figures and images for "The molecular basis for functional divergence of duplicated SOX factors controlling endoderm formation and left-right patterning in zebrafish"

### Supplementary Video 1

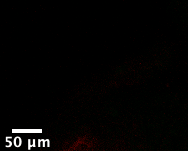
